# Discovering Glycosylation-Dependent Protein Function by Thermal Proteome Profiling

**DOI:** 10.64898/2025.12.06.692499

**Authors:** Johannes F. Hevler, Mirat Sojitra, Tomislav Caval, Michael Schoof, André Mateus, Carolyn R. Bertozzi

## Abstract

Protein glycosylation regulates essential cellular processes including protein folding, stability, and cell-cell interactions; however, how aberrant glycosylation impacts protein function and interaction networks remains poorly understood. Here we combine mass spectrometry-based proteomics, chemical glycobiology, and molecular dynamics simulations to systematically investigate glycosylation dependent protein stability and function. By perturbing the secretory pathway at defined steps, we generated proteins with distinct glycan structures and analyzed their functional consequences using thermal proteome profiling. This approach revealed that cells mount convergent stress responses to diverse glycosylation perturbations, characterized by coordinated trafficking reorganization. This reorganization effectively redirects protein flux from secretion toward degradation. We further identified that perturbing terminal glycan modifications, particularly sialylation and fucosylation, exerts the most profound effects on cell surface protein function. We studied in detail how loss of fucosylation restricts the conformational dynamics of key domains in integrin alpha 4, reducing VCAM-1 binding affinity and suggesting a potential therapeutic strategy for multiple sclerosis. Overall, our systematic approach uncovered extensive complexity in how glycosylation regulates protein function beyond simple glycoform identification and provides a resource for dissecting essential glycan-protein relationships.

## Main

PROTEIN glycosylation is a highly complex co/posttranslational modification that occurs largely within the secretory pathway in eukaryotes (1). Most often attached either to asparagine (*N*-glycosylation) or serine/threonine (*O*-glycosylation) residues, glycans shape protein folding, stability, trafficking, and (glyco)protein-(glyco)protein interactions (2, 3). The specific linkages and branching patterns of these glycans are determined by networks of glycosyltransferases and glycosidases that operate in the ER and Golgi compartments. Because of this structural diversity, glycosylation fine-tunes nearly every aspect of protein behavior from receptor activation to immune recognition, and its dysregulation contributes to cancer, neurodegeneration, and congenital disorders of glycosylation (4–6). Consequently, the identification and characterization of glycosylation patterns of proteins holds great potential for understanding disease mechanisms, and for the development of biomarkers and therapeutic interventions (7–9).

Unfortunately, identifying and characterizing disease-relevant glycosylation patterns remains challenging due to structural complexity and heterogeneity. Despite advances in glycoproteomics, most large-scale studies remain limited to cataloging glycosylation sites and glycoforms rather than linking these modifications to functional consequences in their native cellular context. Traditional approaches often remove glycoproteins from membranes or use recombinant fragments, disrupting the molecular environment where glycans exert their regulatory effects. There is therefore a critical need for integrated strategies that can quantify how specific glycan structures influence protein stability, interactions, and activity within living cells.

Thermal Proteome Profiling (TPP) offers a new approach to glycoproteomics by enabling proteome-wide measurement of protein stability through temperature-dependent solubility changes in the native cellular environment (10). Here, we developed an integrative approach combining TPP and glycoproteomics with chemical glycobiology and molecular dynamics simulations to systematically link glycan structures to protein function. We applied this framework to human myeloid cells treated with small-molecule inhibitors (NGI-1, kifunensine, 3FAX, 2FF) targeting distinct stages of glycan biosynthesis and comprehensively explored the relationship between glycan structures and their functional consequences in living cells. We uncover proteome-wide adaptations to glycosylation stress and identify a core fucosylation–dependent mechanism controlling integrin *α*4*β*1 function, reducing its interaction with the natural ligand VCAM-1 (Vascular Cell Adhesion Molecule-1). Together, these results establish a comprehensive framework for discovering glycosylation-dependent vulnerabilities at proteome scale, with direct implications for cancer, immune regulation, and other diseases characterized by altered glycosylation.

### Global Remodeling of Proteome Stability under Glycosylation Stress

To elucidate the proteome-wide effects of altered protein glycosylation, we first set out to establish a model system that produces glycoproteins with specific glycosylation types. For this, we treated immortalized human myeloid leukemia cells (U937) with small molecule inhibitors (**Fig. 1a**) enabling the targeted inhibition of key enzymes of the glycosylation pathway: oligosaccharyltransferase (OST) complexes (STT3A and STT3B) were inhibited using NGI-1 (*5-(dimethylsulfamoyl)-N-(5-methyl-1,3-thiazol-2-yl)-2-(pyrrolidin-1-yl)benzamide*) to ablate *N*-glycosylation (11); endoplasmic reticulum *α*-1,2-mannosidase I (ERM1) was inhibited using kifunensine (*(5R,6R,7S,8R,8aS)-6,7,8-trihydroxy-5-(hydroxymethyl)-1,5,6,7,8,8a-hexahydroimidazo[1,2-a]pyridine-2,3-dione*) to generate predominantly high-mannose *N*-glycans (12); and peripheral glycosylation of many glycan types were inhibited using 3FAX (13) (*methyl 5-acetamido-2,4-diacetyloxy-3-fluoro-6-[(1S,2R)-1,2,3-triacetyloxypropyl]oxane-2-carboxylate*) and 2FF ((*2S,3R,4R,5S)-2-fluoro-3,4,5-trihydroxyhexanal*), to prevent sialylation (13) and fucosylation (14), respectively (**Fig. 1b**). Flow cytometry using lectin staining (**Fig. 1c, Extended Data Fig. 1, Extended Data Table 1**) and glycoproteomic profiling (**Fig. 1d, Extended Data Table 1**) confirmed the expected glycan alterations, validating the specificity of each perturbation. Under optimized conditions, all treatments achieved strong pathway inhibition with minimal cytotoxicity, enabling proteome-wide analysis with defined glycan alterations. Notably, glycoproteomics revealed that inhibition of sialylation led to an increase in fucosylated glycoproteins. Flow cytometry with *Lotus tetragonolobus* lectin (LTL) confirmed elevated levels of fucose-containing oligosaccharides (**Extended Data Fig. 1, Extended Data Table 1**).

**Fig. 1.**
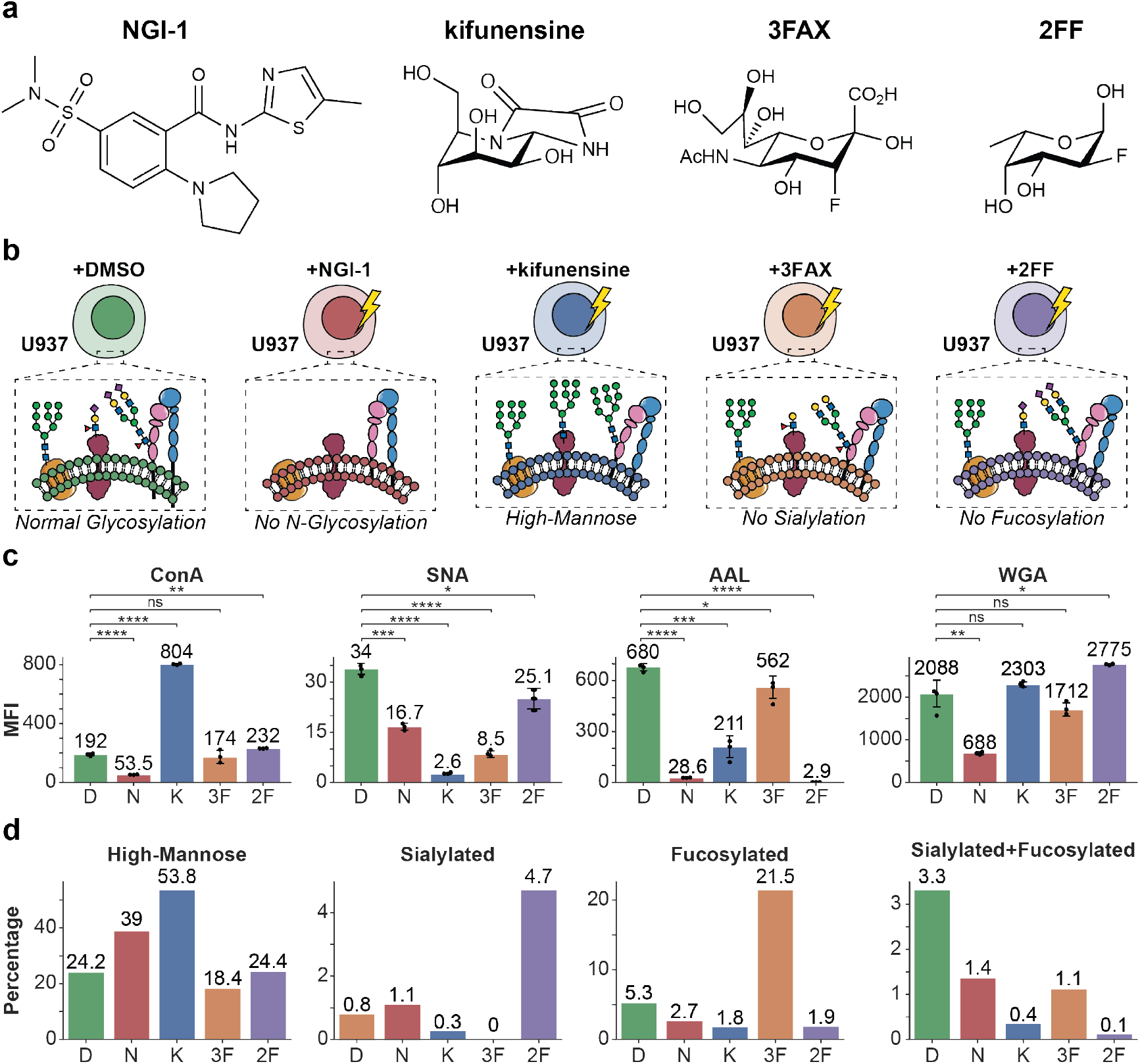
Targeted perturbations of glycan biosynthesis generate proteins with distinct glycan modifications. **(a)** Chemical structures of small molecules used to perturb the glycosylation pathway. **(b)** Experimental strategy for generating distinct glycosylation profiles in U937 cells. Cells were treated with vehicle control (DMSO) or small molecule inhibitors targeting key glycosylation enzymes: NGI-1 (oligosaccharyl transferase complexes), kifunensine (*α*-mannosidase I), 3FAX (sialyltransferases), and 2FF (fucosyltransferases). Schematic diagrams illustrate predicted glycan structures resulting from each treatment. **(c)** Flow cytometric validation of glycan perturbations using fluorescent lectins. Bars show median fluorescence intensity (MFI) for lectins with distinct glycan specificities: Concanavalin-A (ConA, high-mannose binding lectin), *Sambucus nigra* I (SNA, *α*2,6-linked Lac(di)NAc), *Aleuria aurantia* Lectin (AAL, *α*-fucose), and Wheat Germ Agglutinin (WGA, primarily recognizes *N*-acetyl-D-glucosamine and sialic acid containing glycans). Values above bars indicate MFI; individual biological replicates (*n* = 3) shown as dots with error bars representing standard deviation. Statistical significance versus DMSO control determined by unpaired *t*-test with Benjamini-Hochberg correction: ns *p >* 0.05, \**p* ≤ 0.05, \*\**p* ≤ 0.01, \*\*\**p* ≤ 0.001, \*\*\*\**p* ≤ 0.0001. Treatment abbreviations: D (DMSO), N (NGI-1), K (kifunensine), 3F (3FAX), 2F (2FF). **(d)** Glycoproteomic profiling confirms treatment-specific glycan remodeling. Bars show relative percentage (%) of identified glycopeptides bearing high-mannose, sialylated, fucosylated, or both across all treatments: D (DMSO), N (NGI-1), K (kifunensine), 3F (3FAX), 2F (2FF).

We then performed TPP to investigate glycan-dependent changes in protein thermal stability upon treatment with glycosylation inhibitors. TPP leverages the principle that proteins denature and become insoluble upon heat-induced stress (10). By analyzing the remaining soluble fractions at various temperatures using MS, we generated melting profiles for each identified protein. These profiles reflect the specific context of the protein and can be influenced by a variety of factors, including interactions with small molecules (such as drugs or metabolites), nucleic acids, or other proteins, as well as post-translational modifications (15–17). To handle the complexity of melting behaviors and proteome-wide perturbations, we utilized Thermal-Tracks (18), a machine learning framework we previously developed that demonstrates superior performance compared to conventional TPP analysis tools. Using this framework, we systematically evaluated the cellular response to glycosylation pathway disruption and identified proteins that exhibit functional glycan dependencies, including those whose structural integrity, stability, or activity is influenced by specific glycosylation states.

Clustering the melting curves across all conditions revealed eight distinct clusters, each characterized by the melting behavior and melting points (*T*_*m*_). These clusters ranged from proteins with low *T*_*m*_ (53.9 °C, *n* = 2,570) to high *T*_*m*_ (65.6 °C, *n* = 935) (**Extended Data Fig. 2a**), with most proteins melting between 55–57 °C, consistent with previously reported *H. sapiens* meltome data (19). Furthermore, we observed that thermal stability correlates with subcellular localization (**Extended Data Fig. 2b**). Proteins associated with membraneless organelles (e.g., nuclear bodies, nuclear speckles) were enriched among those with *T*_*m*_ values of 53.9–55.3 °C, while cytoplasmic proteins typically melted around 56.5–58.7 °C. In contrast, membrane-embedded proteins (plasma membrane, vesicles) and membrane-bound organelles (endoplasmic reticulum, mitochondria) displayed the highest thermal stability (60.8–65.6 °C), consistent with findings that membrane environments enhance protein thermal stability (19). Following glycosylation pathway perturbation, proteins from this latter group, particularly those residing in the ER, plasma membrane, and vesicles showed an altered melting behavior with decreased thermal stability.

**Fig. 2.**
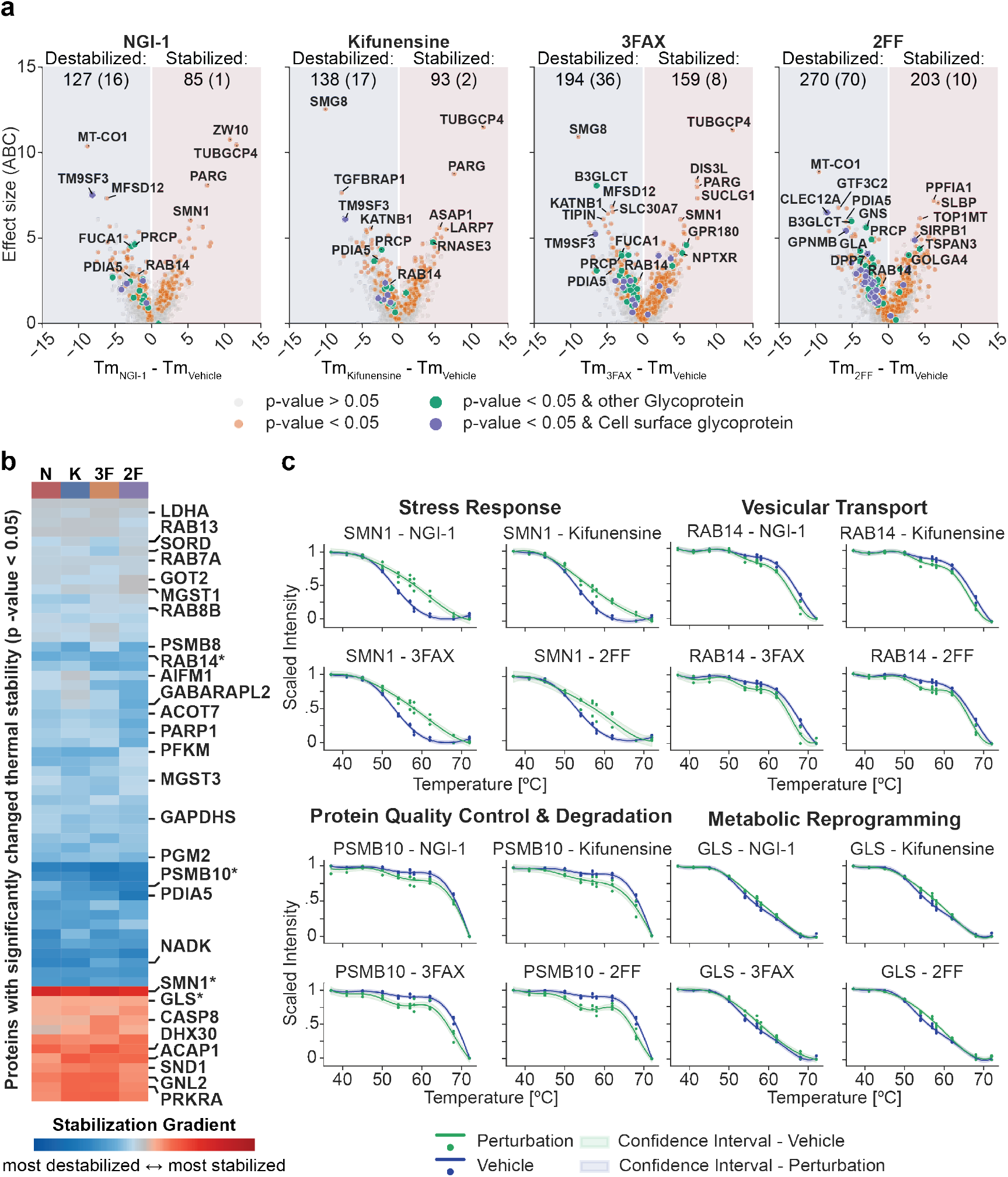
Glycosylation inhibition triggers widespread protein stability changes with convergent cellular responses across multiple pathways. **(a)** Volcano plots showing thermal stability changes (Δ*T*_*m*_) versus effect size (Area between melting curves (ABC)) for proteins following treatment with glycosylation inhibitors NGI-1, kifunensine, 3FAX, and 2FF compared to vehicle control. Proteins are colored by significance (Benjamini-Hochberg adjusted *p*-value *<* 0.05) and classified as glycoproteins (green) or cell surface glycoproteins (purple), other protein (orange) and not significantly changed proteins (gray). Key destabilized and stabilized proteins are labeled. **(b)** Heatmap displaying proteins with significantly changed thermal stability (Benjamini-Hochberg adjusted *p*-value *<* 0.05) across all four glycosylation inhibitor treatments, ordered by similarity of effect size across conditions from destabilized (blue) to stabilized (red). Key proteins are labeled with asterisks (*) indicating proteins whose thermal shift curves are shown in panel c. Treatment abbreviations: N (NGI-1), K (kifunensine), 3F (3FAX), 2F (2FF). **(c)** Thermal shift curves for representative proteins from different functional categories showing dose-dependent stability changes upon glycosylation disruption. Curves show fitted melting curves (lines), confidence intervals (fill) and measured values (circles) for vehicle (blue) and perturbation (green) conditions across the temperature range. Functional categories include stress response, vesicular transport, protein quality control & degradation, and metabolic reprogramming.

To identify specific proteins driving these thermal stability changes, we examined individual protein responses across the four glycosylation inhibitors (NGI-1, kifunensine, 3FAX, 2FF; **Fig. 2**). The inhibitors induced significant thermal stability alterations in 212, 231, 353, and 473 proteins, respectively (**Fig. 2a, Extended Data Table 2**). All inhibitors showed a consistent pattern where more proteins were thermally destabilized than stabilized (~ 60% vs ~ 40%). Perturbations targeting terminal glycan modifications, such as enzymes responsible for adding fucose or sialic acid moieties (2FF and 3FAX), had a more pronounced impact on proteome thermal stability than those targeting early glycosylation steps, such as *N*-glycosylation ablation (NGI-1) or high-mannose *N*-glycan trimming (kifunensine). Among known *N*-glycosylated proteins, sensitivity to 2FF (27%, 80/293) and 3FAX (14%, 44/319) was significantly higher compared to NGI-1 (5%, 17/318) and kifunensine (6%, 19/313). Non-glycosylated proteins were less sensitive to terminal perturbations, affecting 11% (393/3,707) and 8% (309/3,780) by 2FF and 3FAX, respectively. In contrast, early-stage secretory pathway perturbations (NGI-1 and kifunensine) impacted non-glycosylated proteins at similar levels to *N*-glycosylated proteins: 5% (195/3,869) and 6% (212/3,826), respectively (**Extended Data Fig. 2c**). Importantly, the greater thermal destabilization observed with perturbations of terminal glycan modifications (fucosylation and sialylation) compared to early-stage perturbations (*N*-glycosylation ablation or trimming) likely reflects differences in inhibitor potency and cellular tolerance. Inhibitors targeting early glycosylation steps (NGI-1 and kifunensine) exhibit higher cytotoxicity that limits their effective concentration (10 *µ*M for four and seven days, respectively), resulting in partial inhibition and a milder proteomic impact. In contrast, inhibitors of fucosylation and sialylation (2FF and 3FAX) are tolerated at higher doses (100 *µ*M and 50 *µ*M for seven days, respectively), causing more extensive disruption of glycoprotein stability. This difference in achievable inhibitor concentrations likely underlies the stronger proteome destabilization observed with perturbations of terminal glycan modifications.

Across all conditions, secretory pathway perturbation triggered compensatory stabilization of proteins involved in cellular organization, division machinery, and RNA processing (**Extended Data Table 2**). In contrast, GO-enrichment analysis revealed coordinated destabilization of proteins involved in protein folding and ER quality control (e.g., PDIA5), carbohydrate metabolism (e.g., FUCA1), lysosomal degradation (e.g., PRCP), and vesicular transport (e.g., Rab14) (**Fig. 2a, Extended Data Table 2**). Notably, MT-CO1 (mitochondrially encoded cytochrome c oxidase subunit I) was among the most significantly destabilized proteins, consistent with previous observations that ER stress promotes disassembly and degradation of respiratory complex IV (20, 21). Similarly, SMG8, a key nonsense-mediated mRNA decay (NMD) factor, was highly destabilized. Importantly, neither MT-CO1 nor SMG8 are glycoproteins themselves, demonstrating that glycosylation perturbations trigger cascading effects on non-glycosylated proteins through cellular stress response pathways. Since SMG8 negatively regulates SMG1 activity, which initiates NMD, its destabilization would be expected to enhance NMD, representing a compensatory response that suppresses overactivation of the unfolded protein response (UPR) and prevents apoptosis during ER stress (22). These findings align with established stress response mechanisms and validate the biological relevance of our thermal stability measurements.

Despite targeting mechanistically distinct steps, from early glycan attachment to late terminal modifications, the treatments revealed both convergent and unexpected response patterns. A core set of 63 proteins was significantly affected across all four conditions (referred to as pan-responsive proteins, **Extended Data Fig. 2d**). These proteins exhibited exceptionally high concordance in their effect size and altered melting behavior (**Fig. 2b**) regardless of inhibitor mechanism (Pearson correlation 0.91–0.95, **Extended Data Fig. 2d**). However, this tight coordination diminished when examining broader protein sets. Correlations dropped to 0.65–0.81 for proteins affected in at least one condition and further declined to 0.57–0.71 at the proteome-wide level (**Extended Data Fig. 2d**). Notably, inhibitors clustered by their position in the glycosylation pathway: NGI-1 and kifunensine, which disrupt early *N*-glycosylation steps, showed the highest correlations with each other (0.81 for affected proteins, 0.71 proteome-wide), as did 3FAX and 2FF, which both target terminal modifications (0.74 and 0.65, respectively). In contrast, correlations between these two groups were consistently lower. This pattern likely reflects fundamental differences in cellular consequences: early pathway inhibition causes more frequent protein misfolding and ER retention, while terminal modification loss affects surface proteins that are already properly folded. While all correlations remain substantial, indicating a shared glycosylation stress response, these distinct clusters reveal how the timing of glycosylation disruption shapes proteome-wide stability patterns.

The convergent response signature encompassed multiple interconnected systems (**Fig. 2b, Extended Data Table 2**). These included stress response activation (e.g., SMN1, STBD1, PRKRA stabilization; AIFM1, PARP1 destabilization), apoptotic pathway engagement (e.g., CASP8 stabilization), metabolic reprogramming (e.g., GLS stabilization; LDHA, PFKM, GOT2 destabilization), ER quality control disruption (e.g., PDIA5, PSMB8, PSMB10 destabilization), and membrane trafficking reorganization (e.g., multiple Rab GTPases, GABARAPL2 destabilization; ACAP1 stabilization).

The changes in membrane trafficking provide particularly compelling evidence for coordinated cellular reorganization. The thermal destabilization of multiple Rab GTPases alongside the autophagy regulator GABARAPL2, coupled with stabilization of the recycling adaptor ACAP1 (**Fig. 2b**), suggests a systematic breakdown of trafficking systems upon glycosylation inhibition. Rab GTPases likely accumulate in their inactive, GDP-bound cytosolic form, where loss of membrane interactions and effector complexes results in reduced thermal stability (23). Particularly, the destabilization of RAB7A, essential for autophagosome-lysosome fusion, together with GABARAPL2 destabilization, provides converging evidence for impaired autophagy (24).

GABARAPL2 destabilization likely reflects its accumulation in a thermally less stable, non-lipidated cytosolic state rather than its more stable membrane-associated form during active autophagosome maturation (25–27). In contrast, ACAP1 stabilization suggests persistent membrane association, potentially due to recycling arrest or stalled coat turnover. Previous work has shown that overexpressed ACAP1 forms stable complexes with clathrin on endosomal membranes, indicating locked-on coat states that impair cargo recycling (28, 29). Thus, thermal profiling data may reflect similar pathological retention of ACAP1 on membranes during glycosylation inhibition, effectively redirecting cellular trafficking from secretion and recycling toward degradative pathways.

Overall, TPP analysis reveals that glycosylation disruption triggers a unified cellular response characterized by coordinated changes across proteostasis, membrane trafficking, metabolic regulation, and stress signaling. This integrated response demonstrates that cells mount systematic, multipathway adaptations to glycosylation perturbations, with implications for understanding disease pathogenesis and therapeutic strategies targeting the secretory pathway.

### Thermal Co-aggregation Analysis of Key Complexes and Protein-Protein Interactions Uncovers Diverse Cellular Responses to Glycosylation Perturbation

Changes in a protein’s thermal stability can be mediated by several factors, including altered subcellular location, PTMs, and altered interaction profiles. To investigate how glycosylation modulates cellular protein-protein interactions and complex assemblies, we performed thermal proximity coaggregation analysis (TPCA) (30–32). TPCA is based on the principle that interacting proteins co-aggregate upon heat-induced denaturation, resulting in similar melting behaviors among interacting protein pairs and complex members (**Fig. 3a**). To validate that thermal co-aggregation profiles reflect established protein associations, we compared pairwise similarity scores to a curated reference dataset for protein complexes (CORUM). Across all conditions, profiles recovered known binary PPIs and multi-subunit complexes at rates above random expectation, with consistent AUC values between 0.63 and 0.66 (**Extended Data Fig. 3a**), identifying approximately 700 known protein complexes (**Extended Data Fig. 3b**). Spatial Analysis of Functional Enrichment (SAFE) (33) revealed distinct functional clusters, confirming that TPCA captures functionally relevant interactions rather than random associations (**Extended Data Fig. 3c**).

**Fig. 3.**
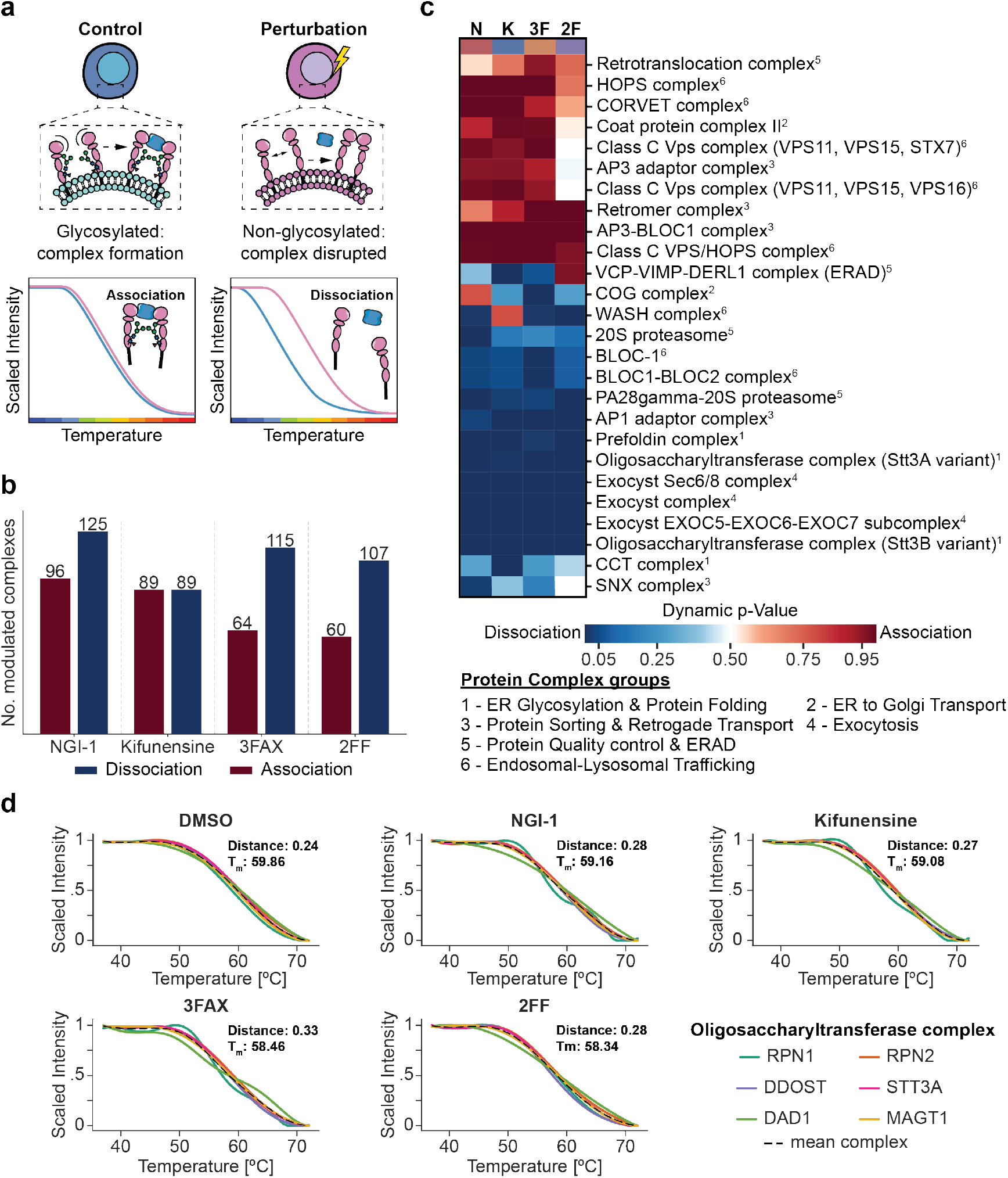
Thermal proximity co-aggregation (TPCA) analysis reveals glycosylation-dependent protein complex remodeling. **(a)** Thermal proximity co-aggregation (TPCA) analysis identifies protein-protein interactions based on correlated thermal aggregation profiles. Proteins in close proximity or forming complexes exhibit similar temperature-dependent aggregation patterns, enabling detection of interaction partners and complex remodeling across conditions. **(b)** Number of protein complexes showing modulated complex behavior compared (blue = dissociation, red = association) upon treatment with glycosylation inhibitors NGI-1, kifunensine, 3FAX, or 2FF compared to DMSO control. **(c)** Heatmap of TPCA profiles for functionally annotated protein complexes across treatments (N: NGI-1, K: kifunensine, 3F: 3FAX, 2F: 2FF). Red indicates convergent profiles (associated complexes), blue indicates divergent profiles (dissociating complexes) relative to vehicle control. Subscript numbers indicate the cellular function of a complex. **(d)** Representative TPCA profiles for the oligosaccharyltransferase complex across treatments. Individual subunit curves (RPN1, RPN2, DDOST, STT3A, DAD1, MAGT1) and mean complex profile (dashed black line) are shown. Correlation of subunits was quantified as the mean pairwise Manhattan distance between fitted melting curves across subunits; Melting temperature (*T*_*m*_) of the complex was defined as the temperature at which normalized abundance reached 0.5, determined by linear interpolation.

To elucidate cellular responses to glycosylation perturbation, we analyzed changes in protein complex modulation following drug induced alteration of glycosylation. When comparing TPCA signatures to vehicle control, we identified slightly more dissociated complexes (90–125 significantly modulated) than associated complexes (60–96 significantly modulated) (**Fig. 3b**). The overall pattern revealed coordinated reorganization of multiple systems: disassembly of translocation machinery (OST complexes), anterograde trafficking complexes (COG and exocyst complexes), proteasomal complexes (20S proteasome complexes) and recycling machinery (WASH complexes), alongside assembly of endolysosomal trafficking complexes (HOPS, CORVET, Class C VPS core, AP3 adaptor, and retromer complexes) (**Fig. 3c**). This suggests a systematic cellular transition from secretion and recycling toward degradative pathways. To better understand the molecular basis of these complex reorganizations, we examined specific protein-protein interactions (PPIs) driving the observed responses. Each treatment extensively rewired the interaction landscape (~ 700–800 altered PPIs per condition; **Extended Data Fig. 3d**), with changes clustering in previously annotated functional modules including RNA processing, transcriptional and translational pathways, vesicle transport, and cell cycle regulation (**Extended Data Fig. 3e**).

The oligosaccharyltransferase (OST) complex provides a particularly informative example of stress-induced reorganization. The OST complex primarily transfers the core Glc_3_Man_9_GlcNAc_2_ glycan to nascent proteins via its STT3A (co-translational) or STT3B (post-translational) variants. However, recent evidence suggests an additional quality control function: Schenkman et al. (34) demonstrated that OST subunits DAD1 and RPN1, which are shared between OST-A and OST-B, can participate in ER-associated degradation pathway (ERAD) by interacting with retrotranslocation machinery. Our data support this dual functionality. We observed OST dissociation not only upon treatment with its direct inhibitor NGI-1, but consistently across all glycosylation perturbations. Differential co-aggregation analysis revealed loss of specific subunit interactions within OST-A (e.g.,STT3A-DDOST) and OST-B (e.g., STT3B-DDOST, STT3B-MAGT1) (**Extended Data Fig. 3d**). Notably, DAD1 and RPN1 showed the most distinct co-aggregation patterns compared to other OST subunits (**Fig. 3d**), providing direct evidence for their delocalization during ER stress. This repurposing of OST subunits toward ERAD could explain the universal OST dissociation and thermal destabilization we observe in cells experiencing prolonged ER stress (**Extended Data Table 3**). The altered co-aggregation patterns of additional OST subunits suggest they too may acquire stress-responsive functions, potentially revealing a broader network of moonlighting proteins that modulate cellular responses to glycosylation stress (35).

Together, these findings demonstrate that glycosylation perturbations trigger coordinated protein complex and interaction network remodeling: dismantling normal quality control and trafficking complexes while forming stress-specific assemblies and reprogramming cellular systems from homeostatic to stress-responsive modes. These adaptive responses reveal the integrated nature of cellular stress mechanisms and potential therapeutic vulnerabilities in glycosylation-altered diseases.

### Alterations in Glycosylation Disrupt Integrin Thermal Stability, Trafficking, and Function

Having established that secretory pathway perturbations induce broad changes in cellular responses, complex formation and protein–protein interactions, we next sought to identify the molecular mechanisms by which altered glycosylation impairs specific protein functions. Among the affected proteins, integrins emerged as particularly noteworthy candidates. This large family of heterodimeric transmembrane receptors, composed of paired *α* and *β* subunits, serves dual critical functions: anchoring cells to extracellular matrix components (including fibronectin, collagen, and VCAM-1) and acting as bidirectional signaling hubs that regulate fundamental cellular processes such as survival, migration, proliferation, and differentiation (36). The functional importance of integrins makes them ideal targets for investigating glycosylation effects, as both subunits are extensively decorated with *N*-linked glycans. These carbohydrate modifications play indispensable roles in multiple aspects of integrin biology, including receptor folding, stability, oligomerization, trafficking, ligand binding, and the conformational transitions between inactive (bent) and active (extended) states of integrins. Notably, specific glycan structures can either promote or inhibit the conformational rearrangements required for integrin activation and productive engagement with the extracellular matrix (37–39).

TPP analysis revealed subtype-specific responses, with *β*1 (ITGB1), *β*2 (ITGB2), *α*4 (ITGA4), and *α*5 (ITGA5) integrins showing differential sensitivity to glycosylation alterations (**Fig. 4a**). The inhibition of fucosylation (2FF) led to strong thermal destabilization of ITGA4 (**Fig. 4a, Extended Data Fig. 4a**), while ITGB1 exhibited significant stabilization upon inhibition of sialylation (3FAX, **Fig. 4a, Extended Data Fig. 4b**). To elucidate whether these thermal stability changes corresponded to alterations in cell surface expression, we performed flow cytometry experiments (**Extended Data Fig. 4c, Extended Data Table 4**). Inhibiting *N*-glycosylation (NGI-1) produced the most dramatic reduction in integrin surface levels, followed by inhibition of mannose trimming (kifunensine), consistent with our hypothesis that early glycosylation disruption causes increased protein misfolding and ER retention. In contrast, terminal glycan perturbations (3FAX and 2FF) had a more modest effect with ITGA4 and ITGB1 even showing increased surface abundance, despite causing significant thermal stability changes. This disconnect suggests that for ITGA4 and ITGB1 terminal glycan modifications may regulate thermal stability through mechanisms independent of trafficking or surface expression.

**Fig. 4.**
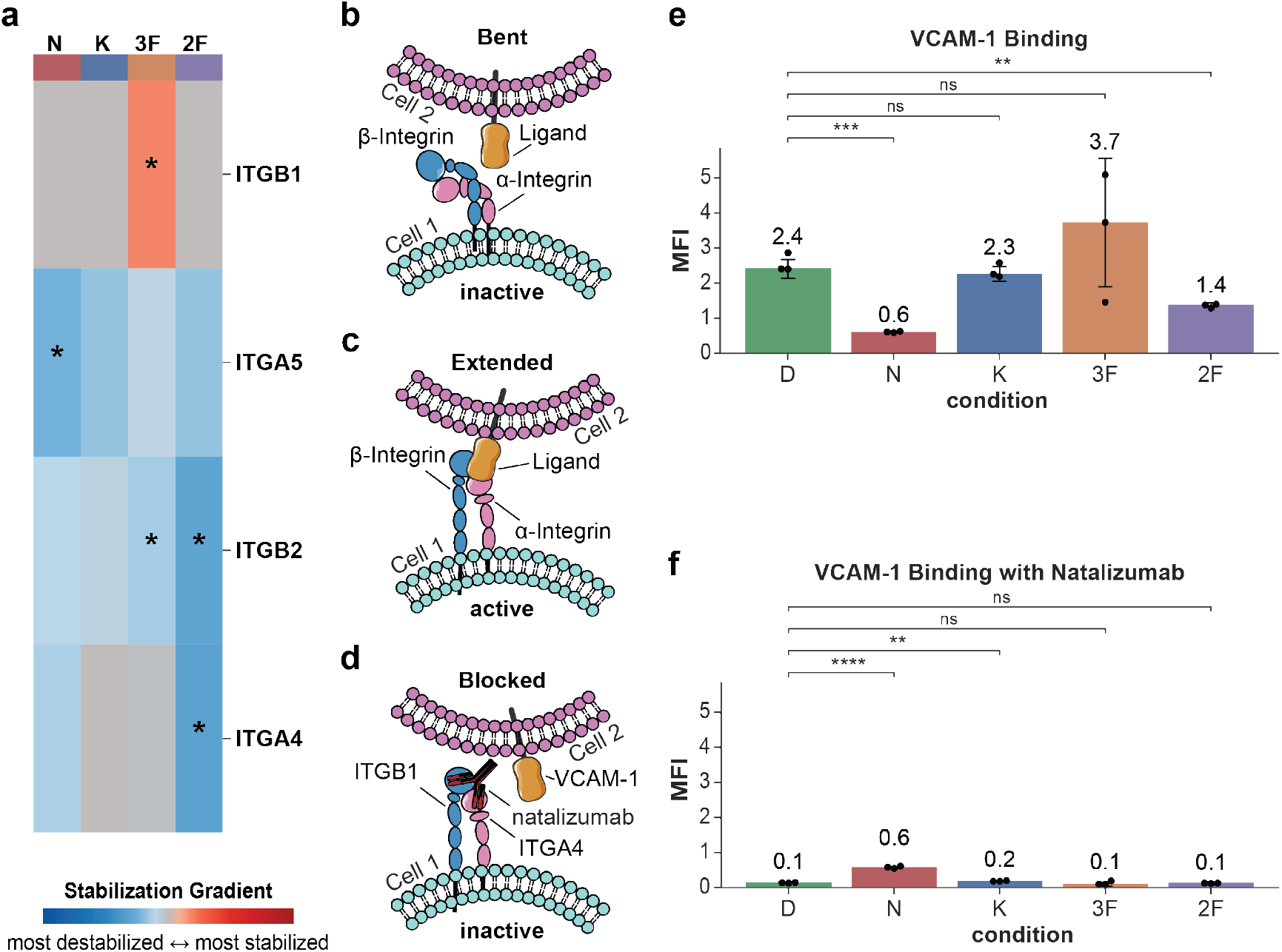
Disruption of the secretory pathway modulates integrin stability, and functional activity. **(a)** Thermal stability analysis of selected integrins showing effect size in stabilization/destabilization upon perturbation with NGI-1 (N), kifunensine (K), 3FAX (3F) and 2FF (2F). Asterisks indicate significantly (Benjamini-Hochberg adjusted *p*-value *<* 0.05) altered thermal stability compared to control (DMSO). Color gradient represents destabilization (blue) to stabilization (red). **(b, c)** Schematic representation of integrin conformational states showing the transition from bent, inactive conformation **(b)** to extended, active conformation **(c). (d)** Schematic representation of VLA-4 heterodimer (*α*4*β*1) which binding to its natural ligand VCAM-1 can be disrupted by the blocking antibody natalizumab. **(e, f)** Functional integrin activity assessed by VCAM-1 binding capacity quantified by mean fluorescence intensity (MFI) under different treatment conditions without **(e)** and with **(f)** addition of the blocking antibody natalizumab. Data represents individual biological replicates (*n* = 3) shown as dots with error bars representing standard deviation. Statistical significance versus DMSO control determined by unpaired *t*-test with Benjamini-Hochberg correction: ns *p >* 0.05, \**p* ≤ 0.05, \*\**p* ≤ 0.01, \*\*\**p* ≤ 0.001, \*\*\*\**p* ≤ 0.0001. Treatment abbreviations: D (DMSO), N (NGI-1), K (kifunensine), 3F (3FAX), 2F (2FF).

The *α*4 and *β*1 subunits form the VLA-4 heterodimer, a principal receptor for VCAM-1 with essential roles in leukocyte trafficking, inflammatory responses, and cancer progression. Like other integrins, VLA-4 undergoes conformational switching between inactive and active states, with only the extended conformation capable of productive VCAM-1 binding (**Fig. 4b–c**). The clinical relevance of VLA-4 is well established, as disruption of the VLA-4-VCAM-1 interaction axis represents an effective therapeutic strategy in multiple sclerosis (40). The monoclonal antibody natalizumab blocks VLA-4-mediated VCAM-1 binding (**Fig. 4d**), thereby preventing pathogenic T-cell migration into the central nervous system. Given this therapeutic precedent, we investigated whether glycosylation-dependent perturbations could alter *α*4*β*1 function and reveal novel vulnerabilities suitable for therapeutic exploitation.

Flow cytometry-based VCAM-1 binding assays revealed that afucosylation (2FF) significantly impaired ligand engagment, beyond the expected effects of *N*-glycosylation inhibition (NGI-1), which drastically reduced VLA-4 surface levels (**Fig. 4e, Extended Data Table 4**). Importantly, afucosylation produced comparable integrin surface levels to kifunensine treatment, yet only afucosylation impaired VCAM-1 binding. This demonstrates that 2FF treatment specifically disrupts integrin function independent of receptor abundance. These flow cytometry results were corroborated by cell adhesion assays, which demonstrated reduced binding of U937 cells to VCAM-1-coated surfaces following afucosylation compared to vehicle control (**Extended Data Fig. 4d**). Conversely, inhibiting the addition of sialic acid (3FAX) resulted in a modest, but non-significant increase in VCAM-1 binding. To confirm that the observed VCAM-1 binding was specifically mediated by *α*4*β*1 integrins, we repeated the binding assays in the presence of the anti-*α*4*β*1 integrin antibody natalizumab. Natalizumab completely abolished VCAM-1 engagement (**Fig. 4f, Extended Data Table 4**), confirming integrin-dependent binding. However, in cells with minimal *N*-glycosylation levels, where *α*4*β*1 surface levels were strongly reduced, addition of natalizumab did not further decrease the already low VCAM-1 binding signal. A likely explanation is that the dramatic loss of surface integrins under *N*-glycosylation inhibition (NGI-1) creates a concentration-dependent limitation, where residual *α*4*β*1 levels are too low for efficient antibody occupancy and complete blocking. Alternatively, it remains possible that inhibiting *N*-glycosylation unmasks integrin-independent interactions with VCAM-1, though this is less likely given the profound reduction in *α*4*β*1 abundance.

The observation that afucosylation reduced VCAM-1 binding despite maintaining high surface levels of both ITGB1 and ITGA4 compared to absence of *N*-glycosylation implicates fucosylation as a direct regulator of *α*4*β*1 activation. This aligns with previous reports highlighting the role of fucosylation in *α*4*β*1 integrin function (41) and suggests that fucose residues are critical for productive ligand engagement. Although not statistically significant in our binding assay, the tendency toward increased VCAM-1 binding following desialylation (3FAX) is consistent with an earlier study demonstrating that monocytes expressing *β*1 subunits lacking *α*2-6-linked sialic acids exhibit enhanced VLA-4-VCAM-1 binding (42). Given our observation of significantly increased fucosylated glycans in cells with impaired sialylation (3FAX) (**Fig. 1b-c, Extended Data Table 1**), we hypothesized that this enhanced binding activity may be partially mediated by elevated fucosylated glycan levels. Specifically, cultured cells treated with 3FAX may exhibit increased (poly-) LacNAc glycan synthesis because inhibited sialyltransferases cannot terminate glycan elongation. These extended glycan structures are subsequently prone to further fucosylation, forming Lewis-type antigens (**Extended Data Fig. 1**) and explaining the marked increase in fucosylated glycans observed with 3FAX treatment (43). This mechanism potentially implicates fucosylated (poly-) LacNAc structures in the enhanced VCAM-1 binding phenotype.

These results demonstrate that distinct glycosylation pathways differentially regulate integrin stability, trafficking, and functional activity. Most importantly, our findings identify fucosylation as a critical determinant of *α*4*β*1–VCAM-1 interactions, revealing a potential vulnerability that can be exploited for therapeutic intervention in diseases characterized by aberrant leukocyte trafficking and inflammation.

### Loss of Core Fucosylation Increases Glycan Flexibility in *α*4-Integrin, Promoting Ca^2+^ Site Shielding and Reducing Conformational Flexibility Through Interactions with the Ca^2+^-Binding Loop and Thigh Domain

Although changes in the *N*-glycosylation profiles of integrins *α*4 and *β*1 are known to impact their functional activity and ligand binding, the underlying molecular mechanisms remains unresolved. To address this, we performed comprehensive glycoproteomics and molecular dynamics (MD) simulations to elucidate how fucosylation controls ITGA4 function at the molecular level.

Our glycoproteomics analysis identified five *N*-glycosylation sites on ITGA4 (N138, N480, N626, N645, and N660; **Extended Data Table 1**). Three sites (N480, N626, N645) displayed consistent glycosylation profiles across all conditions, corroborating previous findings (44). N660 was glycosylated exclusively in kifunensine-treated cells, indicating condition-specific modification. N138 stood out for its glycan heterogeneity beyond high-mannose glycans: in vehicle- and 3FAX-treated cells, we detected fucosylated complex glycans with and without terminal sialic acid (e.g., N4H5F1A1, N4H5F2), whereas 2FF treatment yielded a sialylated complex glycan lacking core fucosylation (N4H5A1; **Extended Data Fig. 5a**). Since high-mannose glycans, which inherently lack core fucose, are present across all conditions including vehicle, the impaired VCAM-1 binding observed specifically upon 2FF treatment indicates that loss of core fucose on mature complex glycans, rather than absence of fucose per se, disrupts integrin function. Importantly, previous glycoproteomic profiling of HEK293 cells showed that surface-enriched ITGA4 proteoforms specifically carry fucosylated complex glycans at N138 (44), directly supporting our finding that core fucose on mature complex glycans is required for integrin function. Further supporting the importance of fucosylation, 3FAX treatment uniquely induced fucosylated poly-LacNAc glycans at N480 and N645, which may underlie the enhanced VCAM-1 binding observed in this condition (**Extended Data Fig. 5b-c**). Our data also suggest that fucosylation is important for ITGB1. Only 3FAX-treated cells displayed a distinctive ITGB1 glycan at N481: a triantennary, desialylated complex structure with core fucosylation, two Lewis-type fucosylations on LacNAc units, and two additional LacNAc repeats (**Extended Data Fig. 5d**).

**Fig. 5.**
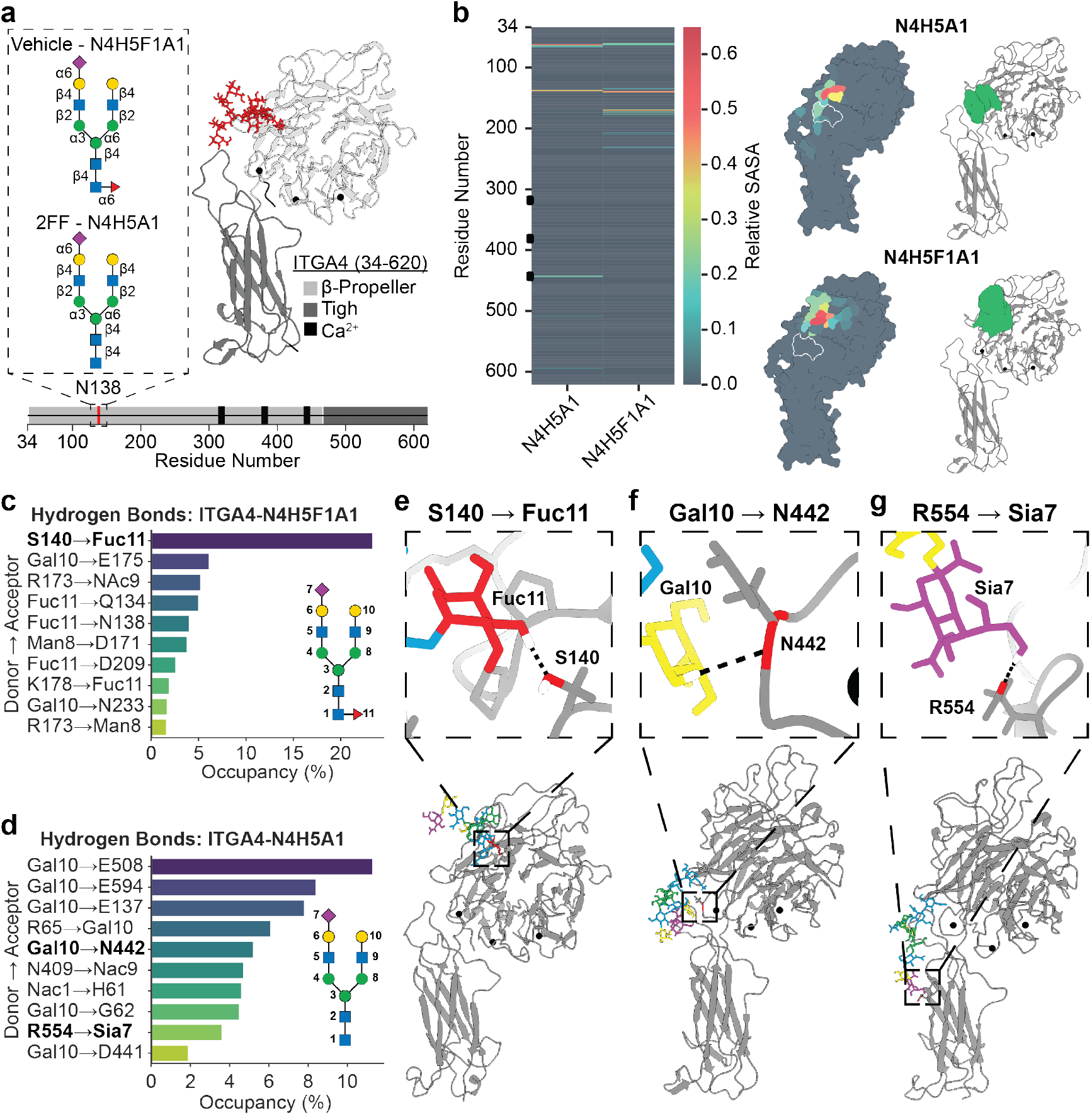
Loss of fucosylation alters ITGA4 glycan dynamics, Ca^2+^-binding site accessibility, and protein flexibility. **(a)** Glycoproteoforms of ITGA4 showing Vehicle-N4H5F1A1 (fucosylated) and 2FF-N4H5A1 (afucosylated) variants with glycan structures mapped onto ITGA4 (residues 34-620) highlighting β-propeller, Thigh, and Ca^2+^-binding domains. For MD simulations, the previously resolved crystal structure of ITGA4 (45) (PDB: 3V4V, chain A) was modified to include missing loops, disulfide bonds, and respective glycosylation. **(b)** Schematic representation of integrin conformational states showing the transition from bent, inactive conformation. **(b)** Comparative glycan shielding analysis demonstrating differential protection of the Ca^2+^-binding region by N4H5A1 (afucosylated) versus N4H5F1A1 (fucosylated) glycans, with heatmap indicating shielding intensity and structural localization. **(c-d)** Hydrogen bond occupancy profiles revealing distinct interaction patterns: **(c)** fucosylated N4H5F1A1 forms contacts restricted to residues near the glycosylation site (N138), whereas **(d)** afucosylated N4H5A1 extends contacts to the Ca^2+^-binding loop (including N442) and Thigh domain. Structural details for H-bond interactions in bold are shown in panel **(e-g). (e)** In fucosylated ITGA4 (N4H5F1A1) core fucose (Fuc11) restricts the conformational phase space of the glycan by hydrogen bonding with S140 near the glycosylation site (N135), **(f)** Galactose (Gal10) interaction with N442 in the Ca^2+^-binding loop, and **(g)** Sialic acid (Sia7) contact with R554 in the Thigh domain. The afucosylated glycan’s increased flexibility enables Ca^2+^-binding site shielding while constraining protein dynamics through frequent glycan-protein contacts, whereas core fucose stabilization in the fucosylated form preserves protein flexibility by limiting distal glycan interactions

To investigate how loss of core fucose affects ITGA4 structure, we performed full-atomistic MD simulations comparing two N138 glycoforms: the fucosylated N4H5F1A1 (abundant in vehicle-treated cells) and the afucosylated N4H5A1 (present in 2FF-treated cells). Simulations used a truncated ITGA4 structure (residues 34–620) encompassing the *β*-propeller, Thigh-, and Ca^2+^-binding domains (**Fig. 5a**). Clustering analysis of torsion angles revealed that core fucosylation rigidifies the glycan backbone, significantly restricting its conformational space compared to the afucosylated glycan (**Extended Data Fig. 6, Extended Data Table 5**). The increased flexibility of the afucosylated glycan resulted in significant reduction of relative solvent accessible surface area (SASA) around the third Ca^2+^ binding site. This shielding effect was confirmed by occupancy analysis over the trajectory, which showed glycan density clearly occluding the Ca^2+^ binding site in the afucosylated form but not in the fucosylated glycan (**Fig. 5b**). A reduced binding of Ca^2+^ also corroborates the observed TPP data for ITGA4 upon 2FF treatment, with apo ITGA4 being more thermally destabilized as compared to the ligand bound form. Together, these findings suggest that loss of core fucosylation impairs Ca^2+^ binding, a crucial step in integrin conformational activation.

Detailed structural analysis of the trajectories further revealed distinct regional flexibility patterns: while the afucosylated glycan itself showed increased flexibility, the ITGA4 backbone exhibited reduced conformational flexibility over the course of the simulation, particularly in the Thigh domain compared to the fucosylated ITGA4 (**Extended Data Fig. 7a-b**). When investigating residue-level flexibility, Thigh domain residues (478–599) were consistently more flexible in fucosylated ITGA4 (ΔRMSF: 1–4 Å). Additionally, especially residues in Ca^2+^-binding loops 2 (residues 377–385) and 3 (residues 439–447) showed greater flexibility in the fucosylated form (ΔRMSF ~ 2 Å) (**Extended Data Fig. 7c-d**).

Contact analyses revealed distinct interaction patterns between the two glycoforms. For the core fucosylated glycan, protein-glycan contacts (distance *<* 5 Å) were limited to residues in the immediate vicinity of the glycosylation site (N138). In contrast, the afucosylated glycan established frequent contacts with residues in the Ca^2+^-binding loop and the Thigh domain, indicating substantially greater spatial reach (**Extended Data Fig. 7e**). Hydrogen bond network analysis identified the molecular basis for these conformational differences (**Fig. 5c-d**). In fucosylated ITGA4, the core fucose (Fuc11) acts as a conformational anchor, restricting glycan mobility through stable hydrogen bonding with residues near the N138 glycosylation site, particularly serine S140 (**Fig. 5e**). Removal of this fucose moiety eliminates these stabilizing interactions, allowing the glycan to adopt extended conformations that form extensive contacts with Ca^2+^-binding loop 3 (residues 439-447) and Thigh domain residues. Specifically, persistent interactions between galactose (Gal10) and N442 (**Fig. 5f**) and between sialic acid (Sia7) and R554 (**Fig. 5g**). These extended conformations physically occlude the Ca^2+^-binding site and restrict movement of functionally critical regions, providing a molecular mechanism for reduced ITGA4 activity upon loss of fucosylation. This reveals a paradox: increased flexibility of afucosylated glycans enables Ca^2+^-binding site occlusion while simultaneously restricting protein dynamics through frequent glycan-protein contacts. Conversely, core fucose anchoring in the fucosylated form preserves protein flexibility by preventing these distal glycan interactions, thereby maintaining the conformational freedom necessary for productive ligand engagement.

To our knowledge, this represents the first molecular-level mechanistic explanation for how fucosylation regulates integrin function. Importantly, our identification of specific glycan-mediated conformational restraints that disrupt ITGA4 function suggests a novel therapeutic strategy: designing small molecules or peptides that mimic the inhibitory contacts made by afucosylated glycans (particularly the Gal10-N442 and Sia7-R554 interactions) could provide an alternative approach to blocking ITGA4 activity. This is particularly relevant for multiple sclerosis treatment, where the current anti-ITGA4/ITGB1 antibody natalizumab, while effective, is restricted to patients without prior John Cunningham virus (JCV) exposure due to the risk of progressive multifocal leukoencephalopathy (PML) (46). An inhibitor that mimics glycan-mediated conformational restriction rather than blocking the binding site entirely could potentially avoid immunosuppression-related complications while maintaining therapeutic efficacy, offering a safer alternative for the broader MS patient population, including those who are JCV positive or have developed neutralizing antibodies to natalizumab.

## Discussion

Understanding the intricate relationship between glycosylation and protein function has important implications for elucidating the molecular basis of diverse diseases and developing targeted therapies. However, due to the extreme diversity in possible glycan structures and modification sites, characterizing these functional dependencies is highly challenging, particularly in a high-throughput, proteome-wide manner. To address this challenge, we employed an integrative approach combining TPP, glycoproteomics, chemical glycobiology, molecular dynamics simulations, and advanced data analysis methods. Central to this strategy was Thermal Tracks, a Gaussian process-based machine learning framework for analyzing TPP data that enables systematic characterization of aberrant glycosylation effects across the proteome with high sensitivity. By integrating glycan structure determination with functional readouts and computational modeling, this multi-faceted approach provides comprehensive insight into how specific glycan modifications regulate protein behavior in native cellular contexts.

It is well established that disrupting the (*N*-)glycosylation biosynthesis pathway has important implications for protein folding, stability, and secretion, and triggers the UPR (47, 48). However, due to the complexity of glycosylation and the UPR, detailed mechanistic insights into how ER stress and the UPR are interconnected with other cellular pathways remain limited. Using TPP in conjunction with Thermal Tracks, we systematically probed the proteome-wide effects of aberrant glycosylation on protein thermal stability and cellular pathway responses. Disrupting glycosylation triggered widespread changes in the thermal stability of proteins across multiple functional categories, including cellular organization, division machinery, stress response, protein degradation, membrane trafficking, and metabolism. These findings reveal extensive crosstalk between the UPR and diverse cellular pathways, providing important new insights into the regulatory networks that respond to glycosylation defects. Understanding these interconnections has significant implications for developing targeted therapeutic strategies for diseases involving, UPR activation and ER stress, such as neurodegenerative disorders and cancer (49).

Glycans can act as conformational switches that stabilize specific protein states, modulate binding interfaces, or alter the accessibility of functional domains (50). Consequently, aberrant glycosylation can shift conformational equilibria toward inactive or pathological states, fundamentally altering protein complexes and PPIs. Using TPCA analysis, we demonstrate that altered *N*-glycosylation induces widespread remodeling of protein complexes. Strikingly, we observed disassembly of critical protein quality control machinery, alongside disruption of secretory and recycling trafficking complexes. Concomitantly, aberrant glycosylation promoted the formation of complexes that redirect misfolded proteins toward direct lysosomal degradation, suggesting a compensatory shift in protein homeostasis mechanisms when canonical quality control pathways are overwhelmed. Beyond quality control machinery, we identified glycosylation-dependent reorganization of the translocation machinery itself, exemplified by stress-induced dissociation of OST complexes and delocalization of DAD1 and RPN1 subunits for repurposing in ERAD pathways. Collectively, our study provides a comprehensive repository of proteins and protein-protein interactions modulated by aberrant glycosylation, offering a valuable resource for understanding glycosylation-dependent regulation of cellular processes and identifying potential therapeutic targets.

While our proteome-wide analysis reveals extensive glycosylation-dependent effects on protein stability and interactions, dissecting the precise molecular mechanisms by which specific glycan modifications regulate individual protein functions remains challenging. Our integrative analysis of ITGA4 provides a paradigm for mechanistically resolving these complex relationships. Although it was previously established that fucosylation of ITGA4 modulates its function (41), the underlying molecular mechanism remained unresolved. By combining TPP as a discovery tool with glycoproteomics and MD simulations, we demonstrate that fucosylation is a critical determinant of the conformational phase space of a *N*-glycan. Loss of core fucosylation dramatically increases glycan flexibility, with profound functional consequences: the more flexible, afucosylated glycans shield the Ca^2+^ binding site of ITGA4 and engage in frequent interactions with the Ca^2+^ binding loop and thigh domain. These glycan-protein interactions effectively restrict the conformational mobility of ITGA4 that is essential for integrin activation (51), providing a molecular explanation for reduced ITGA4 activity in the absence of fucosylation. This mechanistic insight has direct therapeutic implications: mimicking the conformational restraint imposed by afucosylation through rationally designed small molecule inhibitors could provide a novel therapeutic strategy for multiple sclerosis, where ITGA4-mediated leukocyte trafficking drives disease pathology.

This multi-modal strategy, coupled with advanced data analysis frameworks such as Thermal Tracks, enables researchers to move beyond cataloging glycosylation sites to understanding their functional consequences at proteome scale. Our investigation of ITGA4 exemplifies this transition from discovery to mechanistic insight and therapeutic design. Beyond revealing extensive remodeling of protein homeostasis machinery and signaling networks under aberrant glycosylation, our study provides a comprehensive repository of glycosylation-sensitive proteins and interactions to guide future investigations. As the field advances toward precision glycobiology, such integrative approaches will be essential for translating glycosylation insights into targeted therapeutics. By providing both computational tools and a conceptual framework to systematically interrogate glycosylation-dependent regulation, this work enables a new generation of glycobiology research that bridges large-scale discovery with molecular mechanism, ultimately facilitating the development of glycosylation-targeted therapies for diseases ranging from congenital disorders of glycosylation to cancer and autoimmune diseases.

## Material and Methods

### Cell culture and drug treatment

The U937 cells were obtained from ATCC and maintained in RPMI medium supplemented with 10% FBS and 1% penicillin–streptomycin in a T75 flask. The cells were treated with the following inhibitors for 7 days with passaging as needed; i) NGI-1: 10 *µ*M, ii) kifunensine: 10 *µ*M, iii) 3FAX: 50 *µ*M, and iv) 2FF: 100 *µ*M. The vehicle control was treated with DMSO for 7 days. All the cells were cultured at 37 °C at 5% CO_2_.

### Flow-cytometry

#### Lectin or antibody binding to U937 cells

Confluent U937 cells with drug treatment were harvest and ~ 100,000 cells were transferred to V-bottom 96 well plate. The cells in 96 well were centrifuged for 5 min at 400 × g and the supernatant was aspirated. The cell pellet was washed by resuspended it in 200 *µ*L FACS buffer (PBS supplemented with 1% BSA) and centrifuged for 5 min at 400 × g and the supernatant was aspirated. The cell pellet was resuspended in FACS buffer containing fluorophore conjugated lectin at 1 *µ*g/mL and the corresponding inhibitor. For antibody staining, the anti-integrin antibodies were diluted in FACS buffer at 1 in 1000 dilution. The cells were incubated on ice for 1 h and the cells were washed 3 × with FACS buffer. After the final wash, the cell pellet was resuspended in 200 *µ*L FACS buffer containing Sytox Blue dead/alive stain and analyzed using MACSQuant^®^ Analyzer 10 Flow Cytometer.

#### VCAM-1 Binding assay

The VCAM-1 binding assay was done using the same workflow as lectin or antibody binding assay except for following modification. The biotinylated VCAM-1 (Acrobiosystems Cat# VC1-H82F6) was incubated with Streptavidin modified with Alexa647 dye for 1 h on ice in FACS buffer at molar ratio of 4:1 (VCAM-1:Streptavidin) with final concentration of VCAM-1 at 1 *µ*g/mL. This solution was added to pre-washed U937 cells and incubated on ice for 1 h. The cells were washed 3× with FACS buffer and analyzed using MACSQuant^®^ Analyzer 10 Flow Cytometer.

#### Cell adhesion assay

To assess the effect of fucosylation inhibition on cell adhesion, U937 cells that had been pre-treated for 7 days with either DMSO (vehicle control) or 2FF were used for adhesion assays. Twelve-well plates were coated with recombinant human VCAM-1 (0.5 *µ*g/cm^2^) diluted in PBS, or PBS alone as a negative control. Plates were incubated for 2 h at 37°C in 5% CO_2_, followed by overnight incubation at 4°C. Following coating, wells were washed three times with 1 mL PBS per well. Pre-treated cells were harvested, counted, and resuspended in serum-free medium containing their respective treatments (DMSO or 2FF). Cell suspensions containing 5 × 10^5^ cells were added to each well and incubated for 2 h at 37°C in 5% CO_2_ to allow adhesion. After incubation, non-adherent cells were carefully aspirated, and wells were gently washed twice with 1 mL PBS to remove loosely attached cells. Adherent cells were visualized and photographed using bright-field microscopy.

### Sample preparation

#### Thermal proteome profiling

U937 cells were harvested after seven days of drug treatment with either DMSO, NGI-1, kifunensine, 3FAX or 2FF, washed 3 times with cold PBS, and resuspended to a final concentration of 2.5 × 10^8^ cells/mL in PBS. Cell suspensions (20 *µ*L containing 5 × 10^6^ cells) were aliquoted into PCR strip tubes and subjected to a temperature gradient (37.0, 40.9, 44.9, 49.8, 53.9, 56.6, 58.0, 62.0, 68.2, and 72.0°C) for 3 min using a thermal cycler. Following heat treatment, samples were immediately cooled at room temperature for 3. Subsequently, 80 *µ*L of lysis buffer (PBS containing 1% IGEPAL CA-630, 1 mM MgCl_2_, 1 × protease inhibitor cocktail, and 1 U/*µ*L Benzonase) was added to each sample. The PCR plate was incubated for 45 min at 4°C with shaking at 700 rpm to ensure complete cell lysis and nucleic acid digestion. Cell debris and aggregated proteins were pelleted by centrifugation at 2,000 × g for 5 min at 4°C. Soluble protein fractions (70 *µ*L) were transferred to a Multiscreen HTS-HV 96-well filter plate with 0.45 *µ*m PVDF membranes (Merck Millipore) and clarified by centrifugation at 500 × g for 5 min at 4°C. The filtered supernatants were transferred to a new 96-well plate and supplemented with SDS to a final concentration of 2% (w/v) to ensure complete protein denaturation. Samples were stored at − 20°C until further analysis. Proteins were digested according to a modified SP3 protocol (52, 53). In brief, approximately 10 *µ*g of protein from each temperature point (normalized to the sample with the highest concentration (37.0°C)) was diluted to 20 *µ*L with water and transferred to a 96-well plate. Each sample was mixed with 10 *µ*g of carboxylate-modified magnetic beads (Sera-Mag SpeedBeads, Thermo Fisher Scientific; catalog #4515-2105-050250 and #6515-2105-050250, 1:1 mixture) suspended in 10 *µ*L of 15% formic acid and 30 *µ*L of absolute ethanol. Following a 15-min incubation at room temperature with shaking at 500 rpm, protein-bound beads were washed four times with 200 *µ*L of 70% ethanol. On-bead digestion was performed overnight at room temperature by adding 40 *µ*L of digestion buffer containing 100 mM HEPES (pH 8.0), 5 mM chloroacetamide, 1.25 mM TCEP, 200 ng trypsin, and 200 ng Lys-C. Peptides were eluted from the beads using centrifugation, dried under vacuum, and reconstituted in 10 *µ*L of ultrapure water.

For multiplexed quantification, peptides were labeled with TMT10plex reagents (Thermo Fisher Scientific) using the following channel assignments: 126 (37.0°C), 127N (40.9°C), 127C (44.9°C), 128N (49.8°C), 128C (53.9°C), 129N (56.6°C), 129C (58.0°C), 130N (62.0°C), 130C (68.2°C), and 131 (72.0°C). For each reaction, TMT labels (72 *µ*g) were dissolved in 10 *µ*L of anhydrous acetonitrile and added to the corresponding peptide sample. After 1 h incubation at room temperature, the labeling reaction was quenched with 4 *µ*L of 5% hydroxylamine for 15 min. TMT-labeled samples from the same experimental replicate were pooled and acidified with trifluoroacetic acid (TFA) to a final concentration of 1%. Pooled samples were desalted using Waters OASIS HLB *µ*Elution plates (30 *µ*m) according to the manufacturer’s protocol. Briefly, peptides were loaded onto conditioned cartridges, washed with 0.1% TFA, and eluted with 80% acetonitrile containing 0.1% TFA. Eluted peptides were dried under vacuum and subsequently fractionated using high-pH reversed-phase chromatography on C18 spin columns (Thermo Fisher Scientific). Twenty fractions were collected and concatenated by pooling every fifth fraction together, resulting in five final fractions per sample for LC-MS/MS analysis.

#### N-Glyco-proteomics

U937 cells were harvested after seven days of drug treatment and lysed in denaturing buffer containing 7 M urea, 100 mM Tris-HCl (pH 8.5), 1% Triton X-= 100, 50 mM TCEP, 30 mM chloroacetamide (CAA), 2.5 mM MgCl_2_, 1 × protease inhibitor cocktail, and 1 U/*µ*L Benzonase. Lysis was performed for 30 min at 4°C with constant agitation. The pH was adjusted to 8.5 using 10 M NaOH as needed. Cell debris was removed by centrifugation at 18,000 × g for 30 min at 4°C, and the clarified supernatant was collected. Proteins in the supernatant were precipitated using methanol-chloroform precipitation. The resulting protein pellet was air-dried and resuspended in digestion buffer containing 100 mM Tris-HCl (pH 8.5), 1% sodium deoxycholate (SDC), 5 mM TCEP, and 30 mM CAA. Proteolytic digestion was performed by adding trypsin and Lys-C at enzyme-to-substrate ratios of 1:25 and 1:100 (w/w), respectively. Samples were incubated overnight at 37°C with gentle agitation. Following digestion, samples were acidified with trifluoroacetic acid (TFA) and desalted with solid-phase extraction by loading the samples onto a Waters OASIS HLB *µ*Elution Plate (30 *µ*m) and dried under vacuum as described above. Dried peptides were resuspended in glycopeptide enrichment buffer (50 mM sodium carbonate, pH 10.5, in 50% acetonitrile). Phenylboronic acid (PBA)-functionalized silica beads (Bondesil-PBA, 40 *µ*m, Agilent) were used for glycopeptide enrichment at an optimized peptide-to-bead ratio of 1:2.5 (w/w). Prior to use, beads were washed 3–4 times with glycopeptide enrichment buffer. Peptide-bead mixtures were incubated in 96-well plates for 1 h at room temperature with continuous agitation to maintain bead suspension. Following incubation, mixtures were transferred to Multiscreen HTS-HV 96-well filter plates (0.45 *µ*m pore size, Millipore). Non-bound peptides were removed by filtration, and beads were washed seven times with 200 *µ*L of glycopeptide enrichment buffer, using centrifugation at 100 × g for 1 min between washes. Glycopeptides were eluted twice with 50 *µ*L of 1% TFA in 50% acetonitrile, with each elution performed for 15 min at room temperature with shaking at 800 rpm. Eluates were collected in 96-well plates and dried by vacuum centrifugation. Dried glycopeptides were resuspended in SCX loading buffer (5 mM ammonium acetate, pH 2.0) and fractionated using Pierce Strong Cation Exchange spin columns (Thermo Fisher Scientific). Six fractions were collected sequentially: the flow-through from sample loading, followed by step elution’s with 50, 250, 450, 600, and 750 mM ammonium acetate at pH 2.0. All fractions were desalted using Oasis HLB plates as described previously, dried by vacuum centrifugation, and stored at − 20°C until LC-MS/MS analysis.

### LC–MS/MS analysis

#### Thermal proteome profiling

Peptide fractions were reconstituted in loading buffer (0.2% formic acid in MS-grade water) and analyzed using an UltiMate 3000 RSLCnano system (Thermo Fisher Scientific) coupled to an Orbitrap Fusion Tribrid mass spectrometer (Thermo Fisher Scientific). The chromatography system was configured with a trapping cartridge (Acclaim PepMap 100 C18, 5 *µ*m, 100 Å, 300 *µ*m × 20 mm; Thermo Fisher Scientific) and an analytical column (Aurora Series AUR3-25075C18-TS, 25 cm × 75 *µ*m i.d., 1.7 *µ*m C18; IonOptics). Peptides were loaded onto the trapping cartridge at 5 *µ*L/min with solvent A (0.2% formic acid in water) for 5 min, then separated on the analytical column at 300 nL/min using a 95-min gradient. The gradient profile was as follows: 2% solvent B (0.1% formic acid in acetonitrile) for 4 min; 2–4% B over 2 min; 4–28% B over 60.5 min; 28–40% B over 6.5 min; 40–95% B over 5 min; maintained at 95% B for 5 min; followed by reequilibration at 2% B for 12 min. Peptides were ionized using an EASY-Spray ion source operated at 1.85 kV with the column temperature maintained at 45°C and the inlet capillary at 325°C. Data-dependent acquisition (DDA) was performed with the following parameters: MS1 survey scans were acquired from 400–1,450 m/z at 120,000 resolution (at m/z 200) with an automatic gain control (AGC) target of 4 × 10^5^, maximum injection time of 50 ms, and RF lens setting of 60%. The top 10 most intense precursors with charge states between 2+ and 8+ were selected for MS2 fragmentation using a 1.0 m/z isolation window. MS2 scans were acquired in the Orbitrap at 50,000 resolution with an AGC target of 1 × 10^5^, maximum injection time of 120 ms, and normalized HCD collision energy of 38%. The first mass was set to 120 m/z to ensure TMT reporter ion detection. Monoisotopic precursor selection was enabled, and dynamic exclusion was set to 30 s with a *±*10 ppm mass tolerance window.

#### N-Glyco-proteomics

Glycopeptide fractions were reconstituted in loading buffer (0.2% formic acid in MS-grade water) and analyzed using an UltiMate 3000 RSLCnano system (Thermo Fisher Scientific) coupled to an Orbitrap Fusion Tribrid mass spectrometer (Thermo Fisher Scientific). The chromatography system was configured with a trapping cartridge (Acclaim PepMap 100 C18, 5 *µ*m, 100 Å, 300 *µ*m × 20 mm; Thermo Fisher Scientific) and an analytical column (Aurora Series AUR3-25075C18-TS, 25 cm × 75 *µ*m i.d., 1.7 *µ*m C18; IonOptics). Glycopeptides were loaded onto the trapping cartridge at 5 *µ*L/min with solvent A (0.1% formic acid in water) for 5 min, then separated on the analytical column at 300 nL/min using a 120-min gradient optimized for glycopeptide separation. The gradient profile was as follows: 1% solvent B (0.1% formic acid in acetonitrile) for 6.5 min; 5–28% B over 90 min; 28–95% B over 4.5 min; maintained at 95% B for 4 min; followed by re-equilibration at 1% B for 15 min. Peptides were ionized using an EASY-Spray ion source operated at 1.85 kV with the column temperature maintained at 45°C and the inlet capillary at 325°C. Data-dependent acquisition (DDA) was performed with the following parameters: MS1 survey scans were acquired from 500–1,800 m/z at 120,000 resolution (at m/z 200) with an automatic gain control (AGC) target of 4 × 10^5^, maximum injection time of 80 ms, and RF lens setting of 60%. The top 10 most intense precursors with charge states between 2+ and 8+ were selected for MS2 fragmentation using a 2.0 m/z isolation window to accommodate glycopeptide isotopic distributions. MS2 scans were acquired in the Orbitrap at 60,000 resolution with an AGC target of 1 × 10^5^, maximum injection time of 200 ms, and normalized HCD collision energy of 35%. The scan range was set to 120–4,000 m/z to capture both peptide fragments and glycan oxonium ions. Monoisotopic precursor selection was enabled, and dynamic exclusion was set to 30 s with a *±*10 ppm mass tolerance window.

### Data analysis

#### Proteomics search

Raw LC-MS/MS data were processed using two complementary search strategies to maximize peptide and glycopeptide identification. For glycopeptide analysis, pGlyco3 (v3.1) (54) was employed using its *N*-glycosylation HCD fragmentation workflow with the default human *N*-glycan database (pGlyco-N-Human.gdb). For global proteomics and TMT quantification, FragPipe (v20.0) was used, applying the standard TMT10-MS2 workflow (55) of MSFragger (v3.8) (56) for database searching, IonQuant (v1.9.8) for quantification, and Philosopher (v5.0.0) for statistical validation. Prior to FragPipe analysis, raw files were converted to mzML format using MSConvert (ProteoWizard) (57). Both search engines queried the UniProt human reference proteome (UP000005640, downloaded 2023-10-02) supplemented with common contaminant sequences and reversed decoy sequences for false discovery rate (FDR) estimation. Search parameters included: trypsin digestion with up to two missed cleavages; peptide length between 7– 50 (MSFragger) or 7–40 (pGlyco3) amino acids; precursor mass tolerance of 10 ppm; fragment ion mass tolerance of 20 ppm. Fixed modifications included carbamidomethylation of cysteine (+57.021 Da) and TMT10plex labeling of lysine residues and peptide N-termini (+229.163 Da). Variable modifications included oxidation of methionine (+15.995 Da) and protein N-terminal acetylation (+42.011 Da). For MSFragger and pGlyco3 results, peptide-spectrum matches (PSMs) were filtered to 1% FDR using target-decoy competition.

#### Thermal proteome profiling analysis

MSFragger protein-level output files (protein.tsv) were processed using custom Python scripts for quality control, normalization, and scaling prior to thermal stability analysis. Contaminant proteins were removed from the dataset, and proteins lacking gene symbols were assigned to their UniProt accession numbers as identifiers. Temperature points were mapped to their corresponding TMT channels (37, 41, 45, 50, 54, 57, 58, 62, 68, and 72°C for channels T1–T10, respectively). To ensure data quality, proteins with fewer than two quantified unique peptide matches were excluded from further analysis. TMT reporter ion intensities for each temperature point were log_2_-transformed to achieve approximately normal distributions. Infinite values resulting from log-transformation of zero intensities were replaced with NaN and subsequently removed. Median normalization was performed to correct for technical variation between samples. Specifically, for each sample and temperature channel, the median intensity was calculated and adjusted to the global median across all samples. Following normalization, protein intensities were scaled using min-max normalization within each protein and replicate to constrain values between 0 and 1, facilitating comparison across proteins with different absolute abundances. Min and max values were determined per protein per replicate. The processed data were analyzed using Thermal Tracks (18) to compare protein melting behavior between vehicle (DMSO) and glycosylation inhibitor-treated cells (NGI-1, kifunensine, 3FAX, 2FF). For effect-size calculation the area between curves (ABC) was calculated, with the distance of points having overlapping confidence intervals being set to 0. Thermal stability results comparing each compound treatment to vehicle control were visualized using volcano-like plots displaying the melting temperature (*T*_*m*_) difference versus statistical significance. For each protein, *T*_*m*_ was calculated as the temperature at which the remaining soluble fraction reached 50% of the initial value, determined by linear interpolation between adjacent data points. Subsequently, melting curve difference (Δ*T*_*m*_) was calculated as *T*_*m*,vehicle_ − *T*_*m*,drug_, where Δ*T*_*m*_ represents the difference in melting points between protein pairs in each condition. Positive values indicate destabilized proteins (proteins melt faster upon drug perturbation), while negative values represent an increased thermal stability (proteins melt slower upon perturbation). Proteins were annotated based on their glycosylation status and protein subcellular localization using the *N*-GlycositeAtlas (58) and the Cell Surface Protein Atlas (59).

#### Protein melting curve clustering

To identify proteins with similar thermal denaturation profiles, we performed unsupervised clustering of melting curves using Fuzzy C-Means (FCM) clustering. Prior to clustering, proteins were filtered to include only high-quality fits with marginal log-likelihood (MLL) values *>* 0.5 from the Gaussian process regression. Non-melting proteins, defined as those showing minimal thermal-induced aggregation across the temperature range, were excluded from the analysis. Melting curves from both vehicle control and compound-treated conditions were combined, with duplicate control curves removed. FCM clustering was applied using the scikit-fuzzy implementation with the following parameters: fuzziness coefficient (*m*) = 2, convergence tolerance = 1 × 10^−5^, and maximum iterations = 1,000. The optimal number of clusters was determined empirically based on visual inspection of cluster separation and biological interpretability. For each protein and cluster, *T*_*m*_ were calculated as the temperature at which the remaining soluble fraction reached 50% of the initial value, determined by linear interpolation between adjacent data points. Clusters were ordered by their median *T*_*m*_ values for visualization.

#### Gene Ontology Enrichment Analysis

To characterize the subcellular localization and functional properties of proteins within each melting curve cluster, we performed Gene Ontology (GO) term enrichment analysis using the clusterProfiler package (v4.14.6) in R. Protein identifiers from each cluster were converted to Entrez Gene IDs using the org.Hs.eg.db annotation database. Subcellular localization analysis using GO Cellular Component terms was performed exclusively on the melting curve clusters described above. For condition-specific functional characterization, we analyzed proteins showing significant thermal stability changes in response to each compound treatment. Proteins with significantly altered thermal profiles (adjusted *p*-value *<* 0.05) were identified from the thermal stability tracks and categorized into two groups: thermally stabilized and thermally destabilized proteins. Enrichment significance was determined using hypergeometric testing with the entire detected proteome as background. *P*-values were adjusted for multiple testing using the Benjamini-Hochberg method, with an adjusted *p*-value threshold of 0.05 considered significant. Biological Process (BP) GO enrichment analysis was performed separately for stabilized and destabilized protein sets from each treatment condition using the same statistical framework described above. This approach allowed us to identify distinct biological pathways and processes that were differentially affected by compound-induced thermal stabilization versus destabilization, providing insights into the mechanism of action for each glycosylation inhibitor. To identify cluster-specific and shared functional annotations, we performed comparative enrichment analysis across all clusters using the compareCluster function. Results were visualized using enrichplot (v1.26.6) to generate dot plots showing the top enriched terms per cluster, with dot size representing gene ratio and color indicating adjusted *p*-value. Semantic similarity between enriched terms was calculated using DOSE (v4.0.1) to reduce redundancy and identify the most representative biological themes for each cluster. All analyses were performed in R version 4.4 with tidyverse packages for data manipulation.

#### Thermal proximity co-aggregation analysis

To investigate protein complex dynamics and differential protein-protein interactions under glycosylation perturbation, we performed thermal proximity co-aggregation (TPCA) analysis using complementary computational approaches. Protein complex dynamics were assessed using slimTPCA (30), which identifies co-aggregating protein pairs based on correlated melting profiles across the temperature gradient. Differential protein-protein interactions between treatment conditions were analyzed using the R package RTPCA (32), which statistically evaluates changes in co-aggregation patterns between control and perturbed states. The performance of both methods in identifying protein complexes and protein-protein interactions was validated by comparison with experimentally verified complexes from the Comprehensive Resource of Mammalian protein complexes (CORUM) database (60). True positive rates were calculated based on the overlap between predicted co-aggregating pairs and known CORUM complex members. For network-level analysis, protein interaction networks were constructed from significant co-aggregation pairs (FDR < 0.05). Functional modules within these networks were identified using SAFE (Spatial Analysis of Functional Enrichment) (33), implemented through its Python package safepy (https://github.com/baryshnikova-lab/safepy). Identified functional neighborhoods were further manually grouped into broader biological process terms to facilitate interpretation and comparison across treatment conditions. Differential TPCA results comparing each compound treatment to vehicle control were visualized using volcano-like plots displaying melting curve distance versus statistical significance. The melting curve distance metric was calculated as 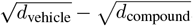, where *d* represents the Euclidean distance between protein pairs in each condition. Positive values indicate gained interactions (proteins co-aggregating more closely upon compound perturbation), while negative values represent lost interactions (proteins co-aggregating less upon treatment). Protein pairs were annotated based on their glycosylation status and protein subcellular localization using the *N*-GlycositeAtlas (58) and the Cell Surface Protein Atlas (59).

#### N-Glyco-proteomics analysis

Because near-isobaric mismatches are a common source of misidentifications for pGlyco3.0, ppmFixer tool (61) was used to determine the most probable glycan topology based on fragment ion patterns for identified *N*-glycopeptides. Glycan compositions were classified into structural categories based on monosaccharide composition using established rules. Specifically, glycans were categorized as: (1) High-Mannose (H ≥ 5, N=2, F=0, A=0), (2) Short *N*-glycans (3<H ≤ 5, N=2, F=0, A=0), (3) Pauci-Mannose (H=3, N=2, F<2, A=0), (4) Hybrid (H ≥ 5, N ≥ 3, H/N ≥ 5/3, with F ≥ 0 or A ≥ 0), (5) Complex (H ≥ 3, N ≥ 4, H/N<5/3, with F ≥ 1 or A ≥ 1), or (6) Unknown for compositions not meeting these criteria. Here, H represents hexose (mannose/galactose), N represents *N*-acetylglucosamine, F represents fucose, and A represents sialic acid residues. For functional analysis, glycans were additionally categorized by their terminal modifications as High-Mannose, Fucosylated (F ≥ 1), Sialylated (A ≥ 1), or Sialylated+Fucosylated (A ≥ 1 and F ≥ 1). To account for glycopeptides containing both sialylation and fucosylation, counts were distributed to both individual categories to accurately represent the prevalence of each modification type. Site-specific glycosylation heterogeneity was visualized using dot matrix plots for proteins of interest. For each glycosylation site and treatment condition, the top *n* most abundant glycoforms were displayed based on summed peak areas for MS1 across replicates. Glycoforms were ordered by structural class (following the hierarchy: High-Mannose, Short *N*-Glycan, Pauci-Mannose, Complex, Hybrid, Unknown) and then alphabetically within each class.

#### Molecular dynamics simulation

##### Input generation for afucosylated and fucosylated ITGA4 simulation systems

The structural model of integrin *α*4 (ITGA4) was constructed based on the crystal structure from PDB entry 3V4V (chain A) (45). This template structure encompasses the extracellular *β*-propeller domain with three bound Ca^2+^ atoms and the thigh domain (residues 34–620), excluding the signal peptide (residues 1–33), transmembrane region, and intracellular domain. A missing loop region in the crystal structure (residues 587–591) was modeled using MODELLER (62) with default loop refinement protocols. Based on our glycoproteomics data revealing differential glycosylation at Asn138 between vehicle- and 2FF-treated cells, we performed in silico glycosylation at this site using the most abundant glycoforms identified experimentally. For vehicle-treated cells, a core-fucosylated complex biantennary glycan was modeled: Neu5Ac(*α*2-6)Gal(*β*1-4)GlcNAc(*β*1-2)Man(*α*1-6)[Gal(*β*1-4)GlcNAc(*β*1-2)Man(*α*1-3)]Man(*β*1-4)GlcNAc(*β*1-4)[Fuc(*α*1-6)]GlcNAc. For 2FF-treated cells, the corresponding afucosylated structure was modeled: Neu5Ac(*α*2-6)Gal(*β*1-4)GlcNAc(*β*1-2)Man(*α*1-6)[Gal(*β*1-4)GlcNAc(*β*1-2)Man(*α*1-3)]Man(*β*1-4)GlcNAc(*β*1-4)GlcNAc.

Glycan structures were built using CHARMM-GUI (63, 64) (www.charmm-gui.org) and disulfide bonds present in the crystal structure were retained to ensure proper structural constraints. The complete models were further processed following the Solution Builder workflow to obtain input files for MD simulations. The glycosylated ITGA4 models were embedded in a rectangular box with 10 nm edge distance from the protein, explicitly solvated with TIP3P water molecules, and neutralized with 150 mM KCl.

##### Molecular dynamics (MD) simulations

All MD simulations were performed with Gromacs (65) (version 2023.1) and CHARM36m all-atom additive forcefields (66) on a High-Performance Computing (HPC) cluster (Sherlock), operated by the Stanford Research Computing Center (https://srcc.stanford.edu/). The system was first energy-minimized using the steepest descent algorithm for 5,000 steps with position restraints applied to backbone atoms (400 kJ/mol/nm^2^) and side chains (40 kJ/mol/nm^2^). Following minimization, the system was equilibrated for 125 ps in the NVT ensemble at 303.15 K using the v-rescale thermostat with a 1 fs integration timestep. Position and dihedral restraints were maintained during equilibration to preserve protein structure and glycan conformations. Production simulations were performed for 100 ns (10 × 10 ns) in the NPT ensemble at 303.15 K and 1 bar, using the v-rescale thermostat (*τ* = 1.0 ps) and C-rescale barostat (*τ* = 5.0 ps) with a 2 fs integration timestep. All bonds to hydrogen atoms were constrained using the LINCS algorithm, allowing the 2 fs timestep. Van der Waals interactions were calculated using a force-switch modifier with a cutoff of 1.2 nm, while electrostatic interactions were computed using the Particle Mesh Ewald (PME) method with a real-space cutoff of 1.2 nm. Coordinates were saved every 100 ps for analysis.

##### Glycan conformational analysis

Glycan conformational dynamics were analyzed using a modified version of the GlycanAnalysisPipeline from GlycoShape (https://github.com/Ojas-Singh/GlycanAnalysisPipeline), as described in Ives et al. (67). The pipeline was adapted to process GROMACS trajectories generated with the CHARMM36m force field. Glycan-only trajectories were extracted from the full MD simulations using the MDAnalysis package in Python, removing protein backbone atoms to isolate glycan conformational behavior. Glycosidic torsion angles (*ϕ, ψ*, and where applicable, *ω*) were calculated for each linkage across the trajectories. Conformational clustering was performed on the torsion angle distributions using the density-based clustering algorithm implemented in the GlycoShape pipeline. The conformational flexibility of each glycoform was quantified by the number of distinct clusters identified, with a higher cluster count indicating greater conformational heterogeneity and flexibility. This approach allows direct comparison of the conformational phase space sampled by fucosylated versus afucosylated glycans, providing insights into how core fucosylation restricts or enhances glycan dynamics at the Asn138 site of ITGA4.

##### Structural analysis of MD simulations

Comprehensive structural analyses were performed using MDAnalysis (68) with custom Python scripts to evaluate the impact of core fucosylation on protein-glycan interactions and dynamics. Root mean square deviation (RMSD) and root mean square fluctuation (RMSF) were calculated for protein backbone to assess structural stability and local flexibility throughout the 100 ns simulations. Protein-glycan interactions were characterized through bidirectional hydrogen bond analysis using geometric criteria (donor-hydrogen distance ≤ 1.2 Å, donor-acceptor distance ≤ 3.0 Å, donor-hydrogen-acceptor angle ≥ 120°), with protein-to-glycan and glycan-to-protein hydrogen bonds analyzed separately to capture all potential interactions. Hydrogen bond occupancy was calculated as the percentage of frames in which specific donor-acceptor residue pairs maintained hydrogen bonds across the trajectory. Additionally, pairwise distance calculations were performed between all protein residues (34–620) and glycan residues for each trajectory frame, computing the minimum distance between atoms of each residue pair to generate comprehensive distance matrices. Both average and minimum distances across the trajectory were analyzed to identify stable and transient protein-glycan contacts, with particular attention to residues 281–289, 344–352, and 406–414 corresponding to Ca^2+^-binding loops in the *β*-propeller domain. Distance values were categorized into interaction zones: strong (<3.5 Å), moderate (3.5–4.5 Å), weak (4.5–5.5 Å), proximal (5.5–10 Å), and distant (*>*10 Å) to quantify how core fucosylation affects the glycan’s spatial sampling and interaction with functional regions of ITGA4.

##### Glycan Shielding and Occupancy analysis

To quantify the extent to which *N*-glycosylation at Asn138 shields the ITGA4 protein surface from solvent exposure, solvent accessible surface area (SASA) calculations were performed using GROMACS with a probe radius of 1.4 Å. SASA values were computed for both the glycosylated protein and the protein alone (with glycan removed) at both residue and atom levels throughout the trajectory. Relative SASA was calculated as (SASA_protein_ – SASA_protein+glycan_) / SASA_protein_, where values approaching 1 indicate complete shielding by the glycan and 0 indicates no shielding effect. This glycan shielding metric was computed for each residue and atom with non-zero SASA in the non-glycosylated state, providing a quantitative map of glycan-mediated surface protection. Mean and maximum shielding values were calculated across the trajectory to identify residues consistently protected by the glycan versus those experiencing transient shielding. Additionally, volumetric occupancy analysis was performed using the VolMap tool in VMD (https://www.ks.uiuc.edu/Research/vmd/) (69, 70) to characterize the three-dimensional space sampled by the glycan throughout the simulation. Occupancy maps were generated with a 1 Å grid resolution, calculating the average probability of glycan atoms occupying each voxel across all trajectory frames. This analysis provided a spatial representation of the glycan conformational ensemble and its coverage of the protein surface.

## Data availability

All raw files, search parameters and search outputs were deposited to the ProteomeXchange Consortium through the PRIDE partner repository with the dataset identifier PXD071666. All MD simulation related data, including the trajectories of the two types of glycosylation discussed here will be deposited to the Zenodo database.

## ACKNOWLEDGEMENTS

This work was supported in part by National Institutes of Health grant CA200423 (C.R.B.), EMBO Postdoctoral Fellowship (ALTF 904-2023; J.F.H.), Howard Hughes Medical Institute Fellowship of the Life Sciences Research Foundation (J.F.H), the Cancer Research Institute Irvington Postdoctoral Fellowship (#CRI5766; M.So.) and Schmidt Science Fellowship, an initiative of Schmidt Futures and the Rhodes Trust (M.Sc.). Some of the computing for this project was performed on the Sherlock cluster. We would like to thank Stanford University and the Stanford Research Computing Center for providing computational resources and support that contributed to these research results.

## AUTHOR CONTRIBUTIONS

Conceptualization and Methodology, J.F.H., A.M. and C.R.B.; Investigation, J.F.H. and M.So.; Verification, J.F.H., M.So., T.C. M.Sc., and A.M.; Formal Analysis and Visualization, J.F.H.; Writing – Original Draft, J.F.H.; Writing – Review & Editing, M.So., T.C., A.M. and C.R.B.; Funding Acquisition, J.F.H., M.So., A.M. and C.R.B.; Resources, C.R.B.; Supervision, C.R.B.

## Bibliography

1. A. Varki et al. Essentials of glycobiology. Cold Spring Harbor Laboratory Press, Cold Spring Harbor, NY, 4th edition, 2022.

2. A. Varki. Biological roles of glycans. Glycobiology, 27:3–49, 2017. doi: 10.1093/glycob/cww086.

3. R. A. Laine. A calculation of all possible oligosaccharide isomers both branched and linear yields 1.05×10(12) structures for a reducing hexasaccharide – the isomer-barrier to development of single-method saccharide sequencing or synthesis systems. Glycobiology, 4: 759–767, 1994. doi: 10.1093/glycob/4.6.759.

4. N. G. Jayaprakash and A. Surolia. Role of glycosylation in nucleating protein folding and stability. Biochemical Journal, 474:2333–2347, 2017. doi: 10.1042/BCJ20170111.

5. Y. van Kooyk and G. A. Rabinovich. Protein-glycan interactions in the control of innate and adaptive immune responses. Nature Immunology, 9:593–601, 2008. doi: 10.1038/ni.f.203.

6. J. Lubbers, E. Rodriguez, and Y. van Kooyk. Modulation of immune tolerance via siglec-sialic acid interactions. Frontiers in Immunology, 9:2807, 2018. doi: 10.3389/fimmu.2018.02807.

7. A. L. Rebelo, M. T. Chevalier, L. Russo, and A. Pandit. Role and therapeutic implications of protein glycosylation in neuroinflammation. Trends in Molecular Medicine, 28:270–289, 2022. doi: 10.1016/j.molmed.2022.01.004.

8. M. H. Hu, Y. Lan, A. Lu, X. X. Ma, and L. J. Zhang. Glycan-based biomarkers for diagnosis of cancers and other diseases: Past, present, and future. Progress in Molecular Biology and Translational Science, 162:1–24, 2019. doi: 10.1016/bs.pmbts.2018.12.002.

9. B. A. H. Smith and C. R. Bertozzi. The clinical impact of glycobiology: targeting selectins, siglecs and mammalian glycans. Nature Reviews Drug Discovery, 20:244, 2021. doi: 10.1038/s41573-021-00160-1. Erratum.

10. M. M. Savitski et al. Tracking cancer drugs in living cells by thermal profiling of the proteome. Science, 346:1255784, 2014. doi: 10.1126/science.1255784.

11. H. Lu, N. A. Cherepanova, R. Gilmore, J. N. Contessa, and M. A. Lehrman. Targeting STT3A-oligosaccharyltransferase with NGI-1 causes herpes simplex virus 1 dysfunction. FASEB Journal, 33:6801–6812, 2019. doi: 10.1096/fj.201802044RR.

12. A. D. Elbein, J. E. Tropea, M. Mitchell, and G. P. Kaushal. Kifunensine, a potent inhibitor of the glycoprotein processing mannosidase-I. Journal of Biological Chemistry, 265:15599–15605, 1990.

13. W. H. D. Bowles and T. M. Gloster. Sialidase and sialyltransferase inhibitors: Targeting pathogenicity and disease. Frontiers in Molecular Biosciences, 8:705133, 2021. doi: 10.3389/fmolb.2021.705133.

14. Y. Zhou et al. Inhibition of fucosylation by 2-fluorofucose suppresses human liver cancer HepG2 cell proliferation and migration as well as tumor formation. Scientific Reports, 7: 11563, 2017. doi: 10.1038/s41598-017-11911-9.

15. A. Mateus et al. Thermal proteome profiling in bacteria: probing protein state. Molecular Systems Biology, 14:e8242, 2018. doi: 10.15252/msb.20188242.

16. A. Mateus et al. The functional proteome landscape of Escherichia coli. Nature, 588:473–478, 2020. doi: 10.1038/s41586-020-3002-5.

17. N. Kurzawa et al. Deep thermal profiling for detection of functional proteoform groups. Nature Chemical Biology, 19:962–971, 2023. doi: 10.1038/s41589-023-01284-8.

18. J. F. Hevler, S. Verma, M. Sojitra, and C. R. Bertozzi. Thermal tracks: A gaussian process-based framework for universal melting curve analysis enabling unconstrained hit identification in thermal proteome profiling experiments. arXiv, 2025. Preprint.

19. A. Jarzab et al. Meltome atlas – thermal proteome stability across the tree of life. Nature Methods, 17:495–503, 2020. doi: 10.1038/s41592-020-0801-4.

20. Z. S. M. Moosavi and D. A. Hood. The unfolded protein response in relation to mitochondrial biogenesis in skeletal muscle cells. American Journal of Physiology-Cell Physiology, 312: C583–C594, 2017. doi: 10.1152/ajpcell.00320.2016.

21. O. Hori et al. Transmission of cell stress from endoplasmic reticulum to mitochondria: enhanced expression of lon protease. Journal of Cell Biology, 157:1151–1160, 2002. doi: 10.1083/jcb.200108103.

22. A. Carreras-Sureda and C. Hetz. Rna metabolism: putting the brake on the UPR. EMBO Reports, 16:545–546, 2015. doi: 10.15252/embr.201540227.

23. A. H. Hutagalung and P. J. Novick. Role of rab GTPases in membrane traffic and cell physiology. Physiological Reviews, 91:119–149, 2011. doi: 10.1152/physrev.00059.2009.

24. L. Zhan et al. Autophagosome maturation mediated by rab7 contributes to neuroprotection of hypoxic preconditioning against global cerebral ischemia in rats. Cell Death and Disease, 8:e2949, 2017. doi: 10.1038/cddis.2017.330.

25. V. Bui et al. Blocking autophagosome closure manifests the roles of mammalian atg8-family proteins in phagophore formation and expansion during nutrient starvation. Autophagy, 21: 1059–1074, 2025. doi: 10.1080/15548627.2024.2443300.

26. S. Ikeda et al. YAP plays a crucial role in the development of cardiomyopathy in lysosomal storage diseases. Journal of Clinical Investigation, 131:e143173, 2021. doi: 10.1172/JCI143173.

27. X. Y. Cao et al. Nascent glycoproteome reveals that N-linked glycosylation inhibitor-1 suppresses expression of glycosylated lysosome-associated membrane protein-2. Frontiers in Molecular Biosciences, 9:899192, 2022. doi: 10.3389/fmolb.2022.899192.

28. J. Dai et al. ACAP1 promotes endocytic recycling by recognizing recycling sorting signals. Developmental Cell, 7:771–776, 2004. doi: 10.1016/j.devcel.2004.10.002.

29. J. Li et al. An ACAP1-containing clathrin coat complex for endocytic recycling. Journal of Cell Biology, 178:453–464, 2007. doi: 10.1083/jcb.200608033.

30. S. Sun et al. Improved in situ characterization of protein complex dynamics at scale with thermal proximity co-aggregation. Nature Communications, 14:7697, 2023. doi: 10.1038/s41467-023-43526-2.

31. C. S. H. Tan et al. Thermal proximity coaggregation for system-wide profiling of protein complex dynamics in cells. Science, 359:1170–1176, 2018. doi: 10.1126/science.aan0346.

32. N. Kurzawa, A. Mateus, and M. M. Savitski. Rtpca: an R package for differential thermal proximity coaggregation analysis. Bioinformatics, 37:431–433, 2021. doi: 10.1093/bioinformatics/btaa682.

33. A. Baryshnikova. Systematic functional annotation and visualization of biological networks. Cell Systems, 2:412–421, 2016. doi: 10.1016/j.cels.2016.04.014.

34. M. Shenkman et al. Oligosaccharyltransferase is involved in targeting to ER-associated degradation. bioRxiv, 2024. Preprint.

35. C. Phoomak et al. The translocon-associated protein (TRAP) complex regulates quality control of N-linked glycosylation during ER stress. Science Advances, 7:eabc6364, 2021. doi: 10.1126/sciadv.abc6364.

36. X. C. Pang et al. Targeting integrin pathways: mechanisms and advances in therapy. Signal Transduction and Targeted Therapy, 8:1, 2023. doi: 10.1038/s41392-022-01259-6.

37. Q. L. Hang et al. N-glycosylation of integrin α5 acts as a switch for EGFR-mediated complex formation of integrin α5β1 to α6β4. Scientific Reports, 6:33507, 2016. doi: 10.1038/srep33507.

38. X. L. Cai, A. M. M. Thinn, Z. L. Wang, H. Shan, and J. Q. Zhu. The importance of Nglycosylation on β3 integrin ligand binding and conformational regulation. Scientific Reports, 7:4656, 2017. doi: 10.1038/s41598-017-04844-w.

39. T. Isaji et al. N-glycosylation of the β-propeller domain of the integrin α5 subunit is essential for α5β1 heterodimerization, expression on the cell surface, and its biological function. Journal of Biological Chemistry, 281:33258–33267, 2006. doi: 10.1074/jbc.M607771200.

40. M. Baiula, S. Spampinato, L. Gentilucci, and A. Tolomelli. Novel ligands targeting α4β1 integrin: Therapeutic applications and perspectives. Frontiers in Chemistry, 7:489, 2019. doi: 10.3389/fchem.2019.00489.

41. W. Li et al. Reduced α4β1 integrin/VCAM-1 interactions lead to impaired pre-B cell repopulation in alpha 1,6-fucosyltransferase deficient mice. Glycobiology, 18:114–124, 2008. doi: 10.1093/glycob/cwm107.

42. A. V. Woodard-Grice, A. C. McBrayer, J. K. Wakefield, Y. Zhuo, and S. L. Bellis. Proteolytic shedding of ST6Gal-I by BACE1 regulates the glycosylation and function of α4β1 integrins. Journal of Biological Chemistry, 283:26364–26373, 2008. doi: 10.1074/jbc.M800836200.

43. C. D. Rillahan et al. Global metabolic inhibitors of sialyl- and fucosyltransferases remodel the glycome. Nature Chemical Biology, 8:661–668, 2012. doi: 10.1038/nchembio.999.

44. C. M. Potel et al. Uncovering protein glycosylation dynamics and heterogeneity using deep quantitative glycoprofiling (DQGlyco). Nature Structural and Molecular Biology, 32, 2025. doi: 10.1038/s41594-025-01485-w.

45. Y. M. Yu et al. Structural specializations of α4β7, an integrin that mediates rolling adhesion. Journal of Cell Biology, 196:131–146, 2012. doi: 10.1083/jcb.201110023.

46. B. Kornek. An update on the use of natalizumab in the treatment of multiple sclerosis: appropriate patient selection and special considerations. Patient Preference and Adherence, 9:675–684, 2015. doi: 10.2147/PPA.S20791.

47. C. Hetz. The unfolded protein response: controlling cell fate decisions under er stress and beyond. Nature Reviews Molecular Cell Biology, 13:89–102, 2012. doi: 10.1038/nrm3270.

48. J. W. Hwang and L. Qi. Quality control in the endoplasmic reticulum: Crosstalk between erad and upr pathways. Trends in Biochemical Sciences, 43:593–605, 2018. doi: 10.1016/j.tibs.2018.06.005.

49. X. Y. Chen, C. R. Shi, M. H. He, S. Q. Xiong, and X. B. Xia. Endoplasmic reticulum stress: molecular mechanism and therapeutic targets. Signal Transduction and Targeted Therapy, 8:352, 2023. doi: 10.1038/s41392-023-01570-w.

50. M. Castelli et al. How aberrant N-glycosylation can alter protein functionality and ligand binding: An atomistic view. Structure, 31:987–1000, 2023. doi: 10.1016/j.str.2023.05.017.

51. S. H. Wang et al. Integrin α4β7 switches its ligand specificity via distinct conformer-specific activation. Journal of Cell Biology, 217:2799–2812, 2018. doi: 10.1083/jcb.201710022.

52. C. S. Hughes et al. Ultrasensitive proteome analysis using paramagnetic bead technology. Molecular Systems Biology, 10:757, 2014. doi: 10.15252/msb.20145625.

53. C. S. Hughes et al. Single-pot, solid-phase-enhanced sample preparation for proteomics experiments. Nature Protocols, 14:68–85, 2019. doi: 10.1038/s41596-018-0082-x.

54. W. F. Zeng, W. Q. Cao, M. Q. Liu, S. M. He, and P. Y. Yang. Precise, fast and comprehensive analysis of intact glycopeptides and modified glycans with pGlyco3. Nature Methods, 18: 1515–1523, 2021. doi: 10.1038/s41592-021-01306-0.

55. S. I. Djomehri et al. Quantitative proteomic landscape of metaplastic breast carcinoma pathological subtypes and their relationship to triple-negative tumors. Nature Communications, 11:1723, 2020. doi: 10.1038/s41467-020-15283-z.

56. A. T. Kong, F. V. Leprevost, D. M. Avtonomov, D. Mellacheruvu, and A. I. Nesvizhskii. MS-Fragger: ultrafast and comprehensive peptide identification in mass spectrometry-based proteomics. Nature Methods, 14:513–520, 2017. doi: 10.1038/nmeth.4256.

57. M. C. Chambers et al. A cross-platform toolkit for mass spectrometry and proteomics. Nature Biotechnology, 30:918–920, 2012. doi: 10.1038/nbt.2377.

58. S. Sun et al. N-GlycositeAtlas: a database resource for mass spectrometry-based human N-linked glycoprotein and glycosylation site mapping. Clinical Proteomics, 16:35, 2019. doi: 10.1186/s12014-019-9254-0.

59. D. Bausch-Fluck et al. A mass spectrometric-derived cell surface protein atlas. PLoS ONE, 10:e0121314, 2015. doi: 10.1371/journal.pone.0121314.

60. G. Tsitsiridis et al. CORUM: the comprehensive resource of mammalian protein complexes 2022. Nucleic Acids Research, 51:D539–D545, 2023. doi: 10.1093/nar/gkac1015.

61. T. M. Adams, P. Zhao, R. Kong, and L. Wells. ppmFixer: a mass error adjustment for pGlyco3.0 to correct near-isobaric mismatches. Glycobiology, 34, 2024. doi: 10.1093/glycob/cwae006.

62. A. Sali. Comparative protein modeling by satisfaction of spatial restraints. Molecular Medicine Today, 1:270–277, 1995. doi: 10.1016/s1357-4310(95)91170-7.

63. S. J. Park et al. CHARMM-GUI glycan modeler for modeling and simulation of carbohydrates and glycoconjugates. Glycobiology, 29:320–331, 2019. doi: 10.1093/glycob/cwz003.

64. S. Jo, T. Kim, V. G. Iyer, and W. Im. CHARMM-GUI: a web-based graphical user interface for CHARMM. Journal of Computational Chemistry, 29:1859–1865, 2008. doi: 10.1002/jcc.20945.

65. D. Van der Spoel et al. GROMACS: Fast, flexible, and free. Journal of Computational Chemistry, 26:1701–1718, 2005. doi: 10.1002/jcc.20291.

66. J. Huang et al. CHARMM36m: an improved force field for folded and intrinsically disordered proteins. Nature Methods, 14:71–73, 2017. doi: 10.1038/nmeth.4067.

67. C. M. Ives et al. Restoring protein glycosylation with GlycoShape. Nature Methods, 21: 2117–2127, 2024. doi: 10.1038/s41592-024-02464-7.

68. N. Michaud-Agrawal, E. J. Denning, T. B. Woolf, and O. Beckstein. MDAnalysis: A toolkit for the analysis of molecular dynamics simulations. Journal of Computational Chemistry, 32: 2319–2327, 2011. doi: 10.1002/jcc.21787.

69. W. Humphrey, A. Dalke, and K. Schulten. VMD: visual molecular dynamics. Journal of Molecular Graphics, 14:33–38, 1996. doi: 10.1016/0263-7855(96)00018-5.

70. J. Cohen, A. Arkhipov, R. Braun, and K. Schulten. Imaging the migration pathways for O_2_, CO, NO, and Xe inside myoglobin. Biophysical Journal, 91:1844–1857, 2006. doi: 10.1529/biophysj.106.085746.

