## Supplementary Figures for "Discovering Glycosylation-Dependent Protein Function by Thermal Proteome Profiling"

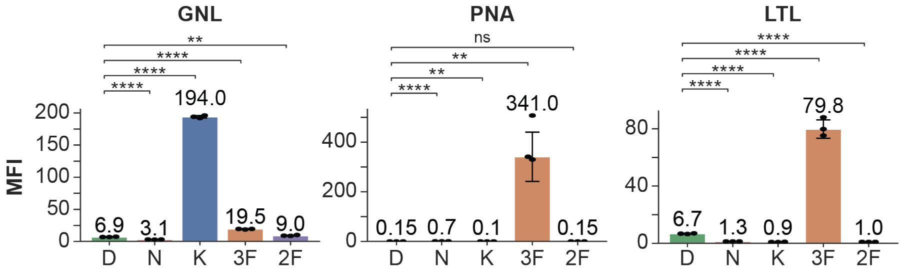


**Extended Data Fig. 1. Targeted perturbations of glycan biosynthesis generate proteins with distinct glycan modifications**. Flow cytometric validation of glycan perturbations using fluorescent lectins. Bars show median fluorescence intensity (MFI) for lectins with distinct glycan specificities: *Galanthus nivalis* lectin (GNL, high-mannose binding lectin), Peanut Agglutinin (PNA, T-antigen binder with preferential binding for terminal Galβ1,3GalNAc), *Lotus tetragonolobus* lectin (LTL, fucose binding with Lewis$^{x}$ as its main recognition motif). Values above bars indicate MFI; individual biological replicates (n=3) shown as dots with error bars representing standard deviation. Statistical significance versus DMSO control determined by unpaired t-test with Benjamini-Hochberg correction: ns $p>0.05$, *$p\leq0.05$, **$p\leq0.01$, ***$p\leq0.001$, ****$p\leq0.0001$. Treatment abbreviations: D (DMSO), N (NGI-1), K (Kifunensine), 3F (3FAX), 2F (2FF).

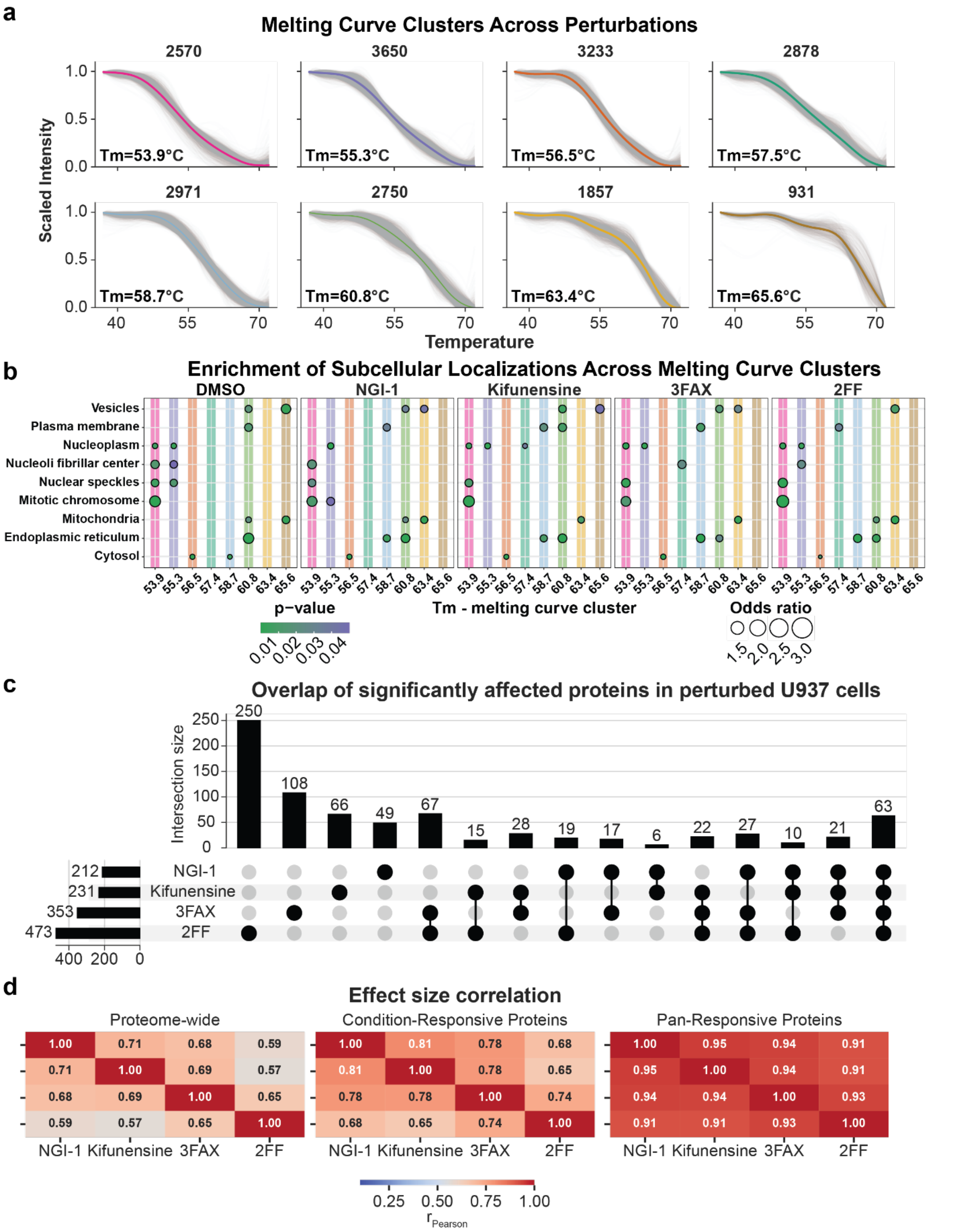


**Extended Data Fig. 2. Melting curve clustering reveals treatment-specific proteomic responses and subcellular organization. (a)** Representative melting curve clusters identified across all perturbations. Proteins are grouped by similar thermal stability profiles (melting curves). Numbers above each panel indicate the number of proteins in each cluster; Tm indicates the average melting temperature. Shaded areas represent individual protein melting curves around the mean cluster profile (bold line). Colors distinguish different cluster profiles. **(b)** Enrichment of subcellular localizations across melting curve clusters for each treatment (DMSO, NGI-1, Kifunensine, 3FAX, 2FF). Dot color intensity indicates enrichment p-value, dot size represents odds ratio. Clusters are ordered by melting temperature (Tm) as shown on the x-axis. **(c)** Overlap of significantly (adj. p-value < 0.05) affected proteins across glycosylation inhibitor treatments. UpSet plot shows intersection sizes between treatments. Numbers above bars indicate the size of each intersection. Horizontal bars show total significantly affected proteins per treatment: 2FF (475), 3FAX (353), Kifunensine (231), NGI-1 (212). **(d)** Effect size correlation matrices showing Pearson correlation coefficients (r) between treatments. Left: proteome-wide correlations across all quantified proteins. Middle: condition-responsive proteins (significantly affected by at least one treatment). Right: pan-responsive proteins (affected by all treatments). Higher correlations in pan-responsive proteins indicate consistent directional changes across perturbations for commonly affected proteins.

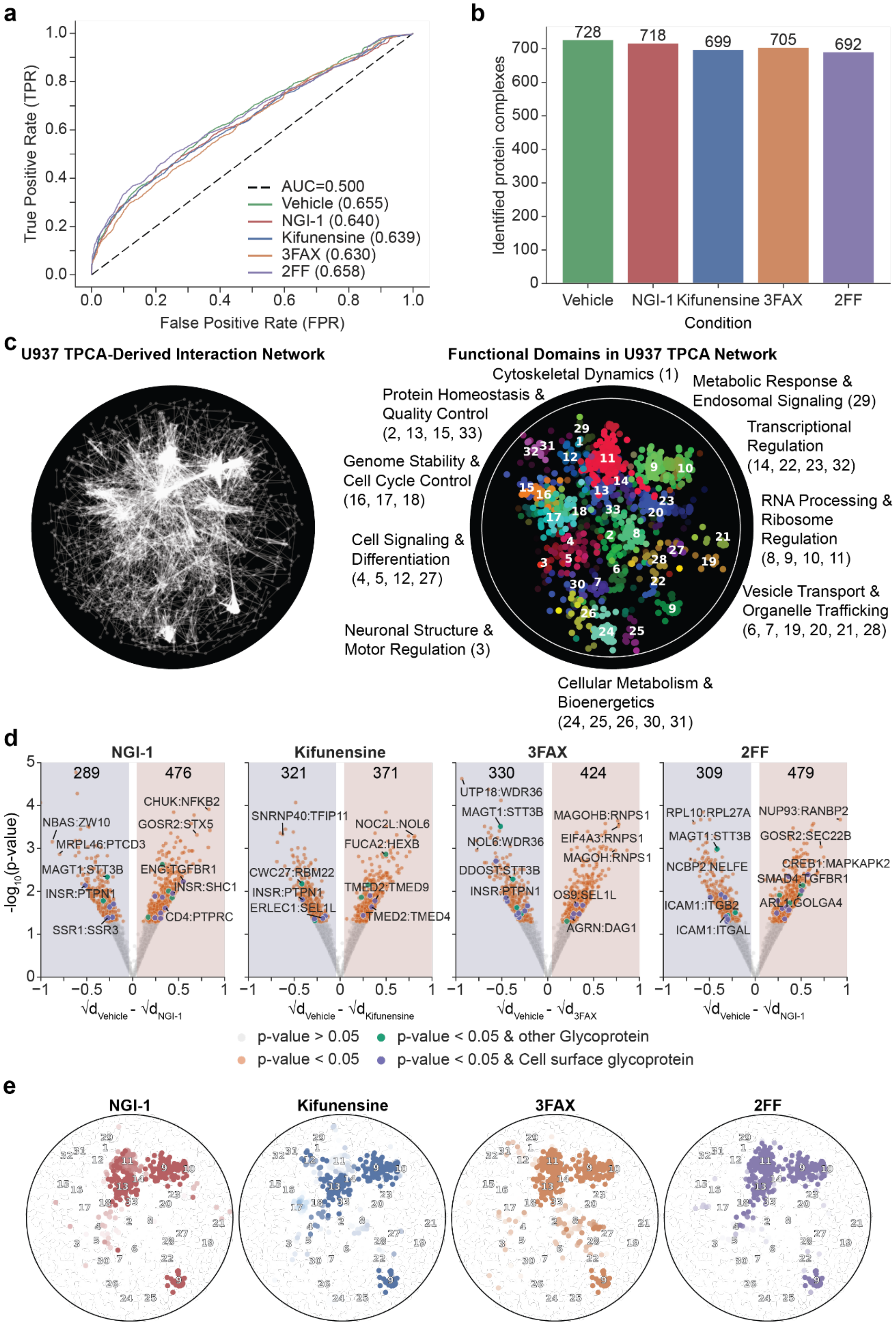
a
AUC=0.500
Vehicle (0.655)
NGI-1 (0.640)
Kifunensine (0.639)
3FAX (0.630)
2FF (0.658)
True Positive Rate (TPR)
False Positive Rate (FPR)
b
728
718
699
705
692
Identified protein complexes
Vehicle
NGI-1
Kifunensine
3FAX
2FF
Condition
c
U937 TPCA-Derived Interaction Network
Functional Domains in U937 TPCA Network
Cytoskeletal Dynamics (1)
Metabolic Response & Endosomal Signaling (29)
Protein Homeostasis & Quality Control (2, 13, 15, 33)
Transcriptional Regulation (14, 22, 23, 32)
Genome Stability & Cell Cycle Control (16, 17, 18)
RNA Processing & Ribosome Regulation (8, 9, 10, 11)
Cell Signaling & Differentiation (4, 5, 12, 27)
Vesicle Transport & Organelle Trafficking (6, 7, 19, 20, 21, 28)
Neuronal Structure & Motor Regulation (3)
Cellular Metabolism & Bioenergetics (24, 25, 26, 30, 31)
d
NGI-1
Kifunensine
3FAX
2FF
-log10(p-value)
289
476
321
371
330
424
309
479
NBAS:ZW10
MRPL46:PTCD3
MAGT1:STT3B
INSR:PTPN1
SSR1:SSR3
CHUK:NFKB2
GOSR2:STX5
ENG:TGFBR1
INSR:SHC1
CD4:PTPRC
SNRNP40:TFIP11
CWC27:RBM22
INSR:PTPN1
ERLEC1:SEL1L
NOC2L:NOL6
FUCA2:HEXB
TMED2:TMED9
TMED2:TMED4
UTP18:WDR36
MAGT1:STT3B
NOL6:WDR36
DDOST:STT3B
INSR:PTPN1
MAGOHB:RNPS1
EIF4A3:RNPS1
MAGOH:RNPS1
OS9:SEL1L
AGRN:DAG1
RPL10:RPL27A
MAGT1:STT3B
NCBP2:NELFE
ICAM1:ITGB2
ICAM1:ITGAL
NUP93:RANBP2
GOSR2:SEC22B
CREB1:MAPKAPK2
SMAD4:TGFBR1
ARL1:GOLGA4
√dVehicle - √dNGI-1
√dVehicle - √dKifunensine
√dVehicle - √d3FAX
√dVehicle - √dNGI-1
p-value > 0.05
p-value < 0.05
p-value < 0.05 & other Glycoprotein
p-value < 0.05 & Cell surface glycoprotein
e
NGI-1
Kifunensine
3FAX
2FF

**Extended Data Fig. 3. Thermal proximity co-aggregation analysis (TPCA) for identifying modulated protein complexes and protein-protein interactions (a)** ROC analysis to quantify the predictive power of TPCA to identify known protein-protein interactions from the CORUM database in U937 cells. Area under the curve (AUC) values are shown for vehicle control (0.655) and glycosylation inhibitor treatments (NGI-1: 0.640, Kifunensine: 0.639, 3FAX: 0.630, 2FF: 0.658), all exceeding random prediction (AUC = 0.500, dashed line). **(b)** Number of TPCA-identified protein complexes in U937 cells across control and glycosylation inhibitor conditions, showing robust complex detection despite perturbations (vehicle: 728, NGI-1: 718, Kifunensine: 699, 3FAX: 705, 2FF: 692 complexes). **(c)** TPCA reveals functional interactome organization in U937 cells. Left: Complete TPCA-derived protein-protein interaction network. Right: Spatial Analysis of Functional Enrichment (SAFE) revealed functional clusters, confirming that TPCA captures functionally relevant interactions. Clusters are colored by biological process with cluster numbers shown in parentheses representing unique GO terms that are further grouped into corresponding biological processes. **(d)** Differential TPCA analysis identifies modulated protein-protein interactions in response to glycosylation inhibition. Volcano plots show change in average Euclidean melting curve distance (Δd) of protein pairs in control (vehicle) or perturbation. Orange: significantly changed PPIs (p-value < 0.05); green: significantly changed PPI including a glycoprotein; purple: significantly changed PPI including a cell surface glycoprotein; gray: non-significant (p-value > 0.05). Numbers indicate proteins with significant TPCA changes per treatment. **(e)** Affected PPIs in the tested perturbations visualized on the U937 cell TPCA interaction network shown in panel c, indicating treatment-specific TPCA changes across functional domains (numbered clusters from panel c).

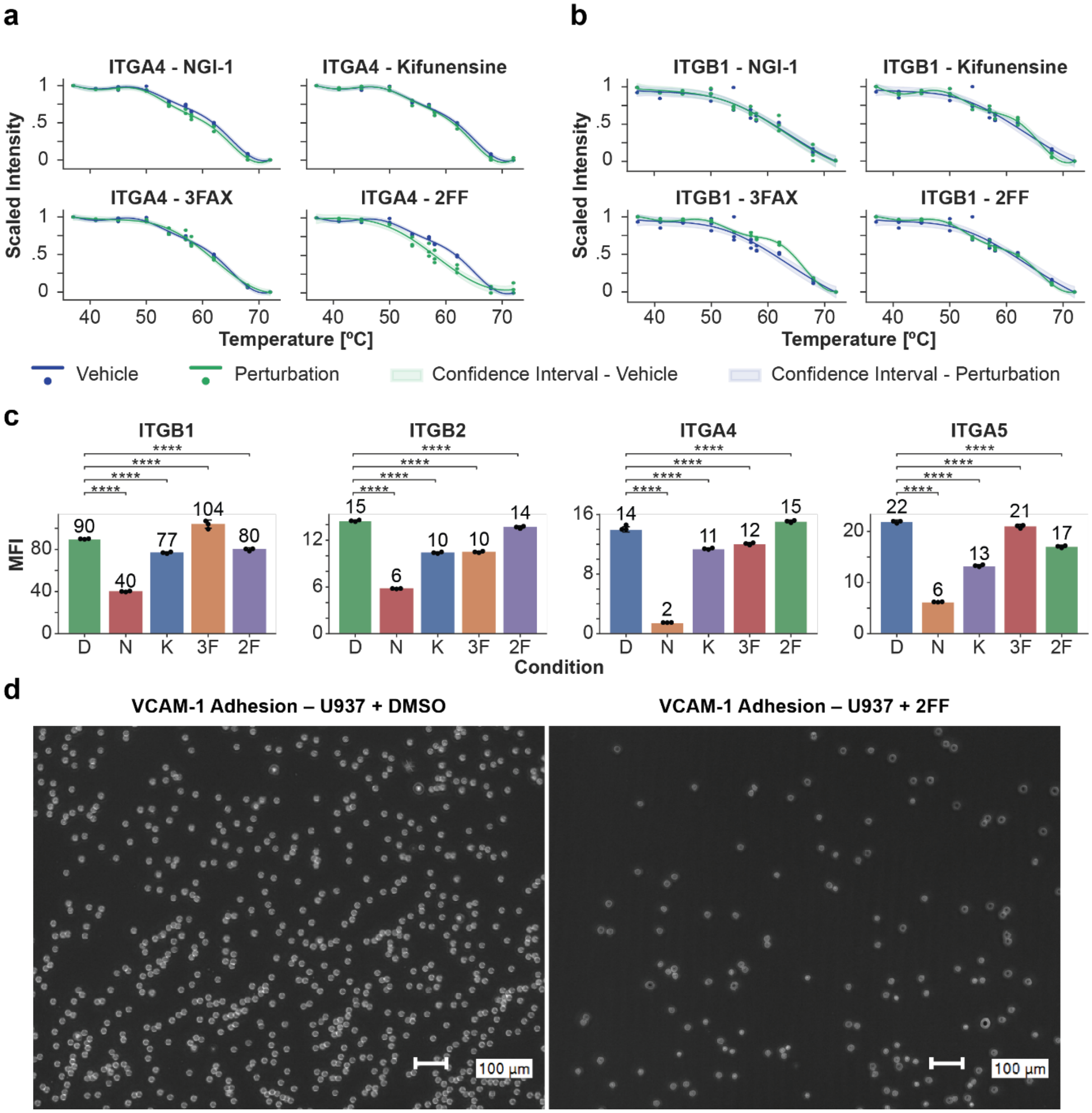


**Extended Data Fig. 4. (a)** ITGA4 melting behavior across glycosylation inhibitor treatments. Thermal stability curves comparing vehicle control (blue) and perturbation conditions (green) for NGI-1, Kifunensine, 3FAX, and 2FF treatments. Shaded areas represent confidence intervals. **(b)** ITGB1 melting behavior across glycosylation inhibitor treatments. Thermal stability curves comparing vehicle control (blue) and perturbation conditions (green) for NGI-1, Kifunensine, 3FAX, and 2FF treatments. Shaded areas represent confidence intervals. **(c)** Surface expression levels of ITGB1, ITGB2, ITGA4, and ITGA5 quantified by mean fluorescence intensity (MFI) following treatment with vehicle control (DMSO (D)) and NGI-1 (N), Kifunensine (K), 3FAX (3F), 2FF (2F). Numbers above bars indicate MFI values. Data represent biological triplicates (black dots), with error bars indicating standard. Statistical significance versus DMSO control determined by unpaired t-test with Benjamini-Hochberg correction: ns $p>0.05$, *$p\leq0.05$, **$p\leq0.01$, ***$p\leq0.001$, ****$p\leq0.0001$. **(d)** Functional validation of integrin-mediated cell adhesion. Representative fluorescence microscopy images of U937 cell adhesion to VCAM-1-coated surfaces following DMSO (left) or 2FF (right) treatment. 2FF treatment substantially reduces cell adhesion, consistent with altered integrin function. Scale bars: 100 µm.

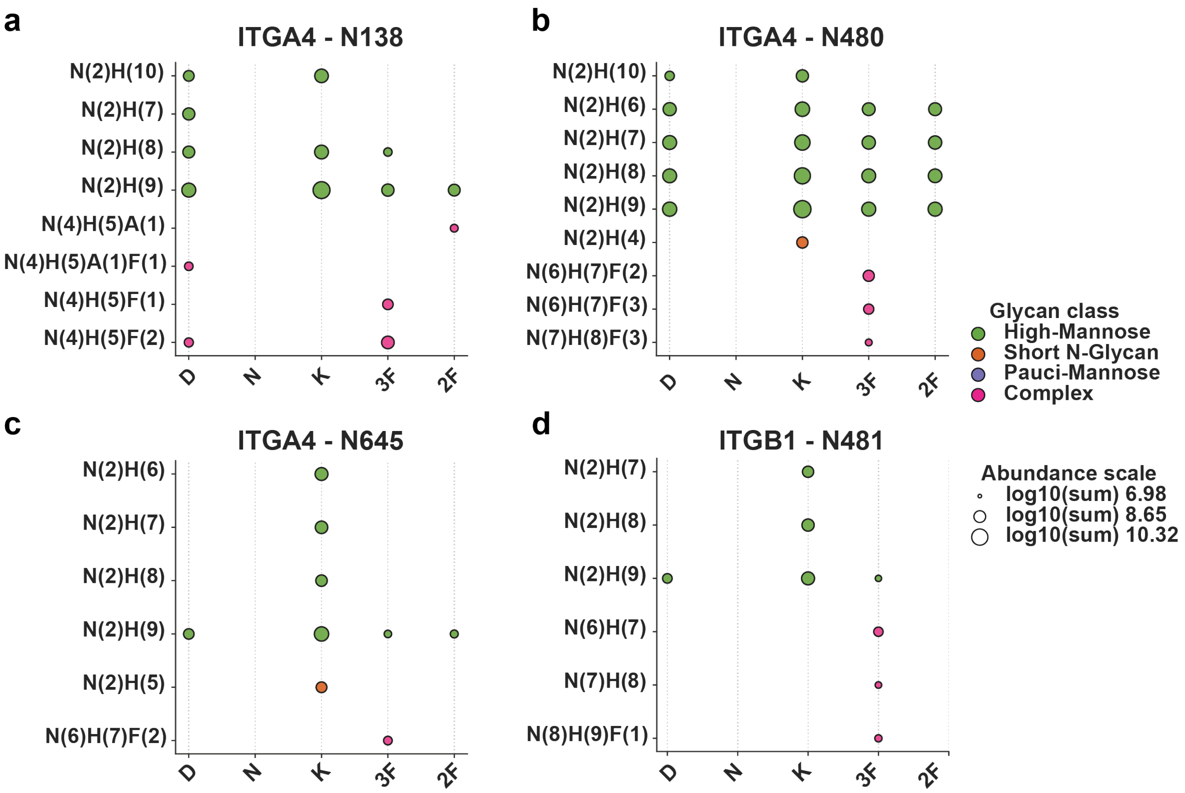


**Extended Data Fig. 5. Site-specific glycoproteomics reveals treatment-dependent glycan remodeling on integrins (a-d). (a)** Glycan composition at ITGA4 N138 across treatments. Dot plots show detected glycan structures with N-glycan composition notation (N = HexNAc, H = Hexose, F = Fucose). **(b)** Glycan composition at ITGA4 N480 across treatments. **(c)** Glycan composition at ITGA4 N645. **(d)** Glycan composition at ITGB1 N481. For all panels, dot size indicates relative abundance on a log10 scale. Dot color represents glycan class: green = high-mannose, orange = short N-glycan, purple = pauci-mannose, pink = complex glycans. Treatment conditions shown: DMSO (D), NGI-1 (N), Kifunensine (K), 3FAX (3F), 2FF (2F).

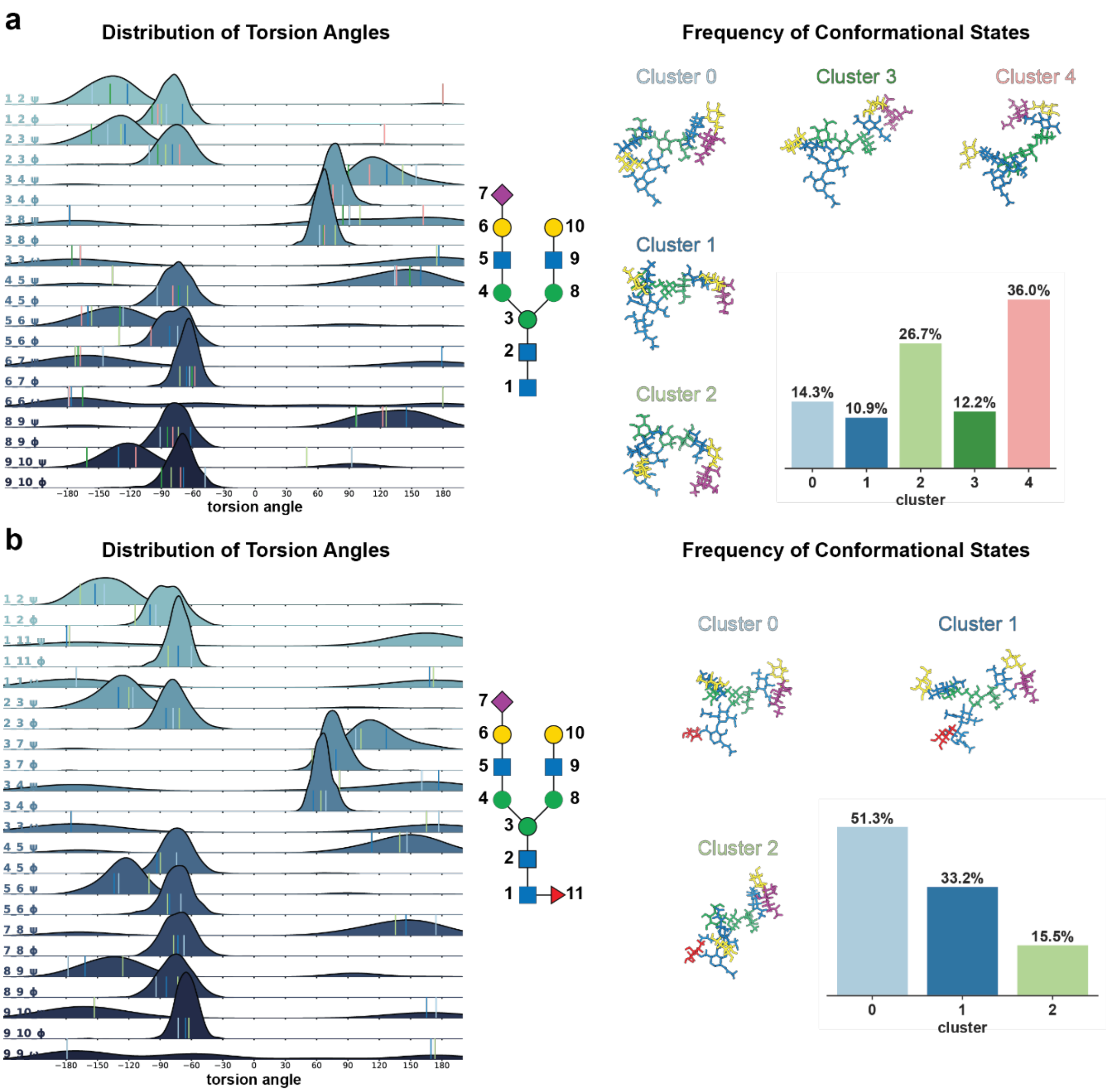


**Extended Data Fig. 6. Loss of core fucosylation increases glycan conformational flexibility and alters the accessible conformational landscape (a-b). (a)** Afucosylated ITGA4 N138 glycan conformational analysis. Left: Distribution of torsion angles (ϕ, ψ, ω) for each glycosidic linkage (numbered 1-10) throughout the MD simulation. Right: Conformational clustering reveals five major glycan conformational states (clusters 0-4) with representative structures shown. **(b)** Fucosylated ITGA4 N138 glycan conformational analysis. Left: Distribution of torsion angles (ϕ, ψ, ω) for each glycosidic linkage (numbered 1-11) throughout the MD simulation. Right: Conformational clustering reveals three major states (clusters 0-2) with representative structures. Glycan moieties are colored according to the SNFG color scheme, with GlcNAc being blue, Man being green, Gal being yellow, NeuAc being violet and Fuc being red.

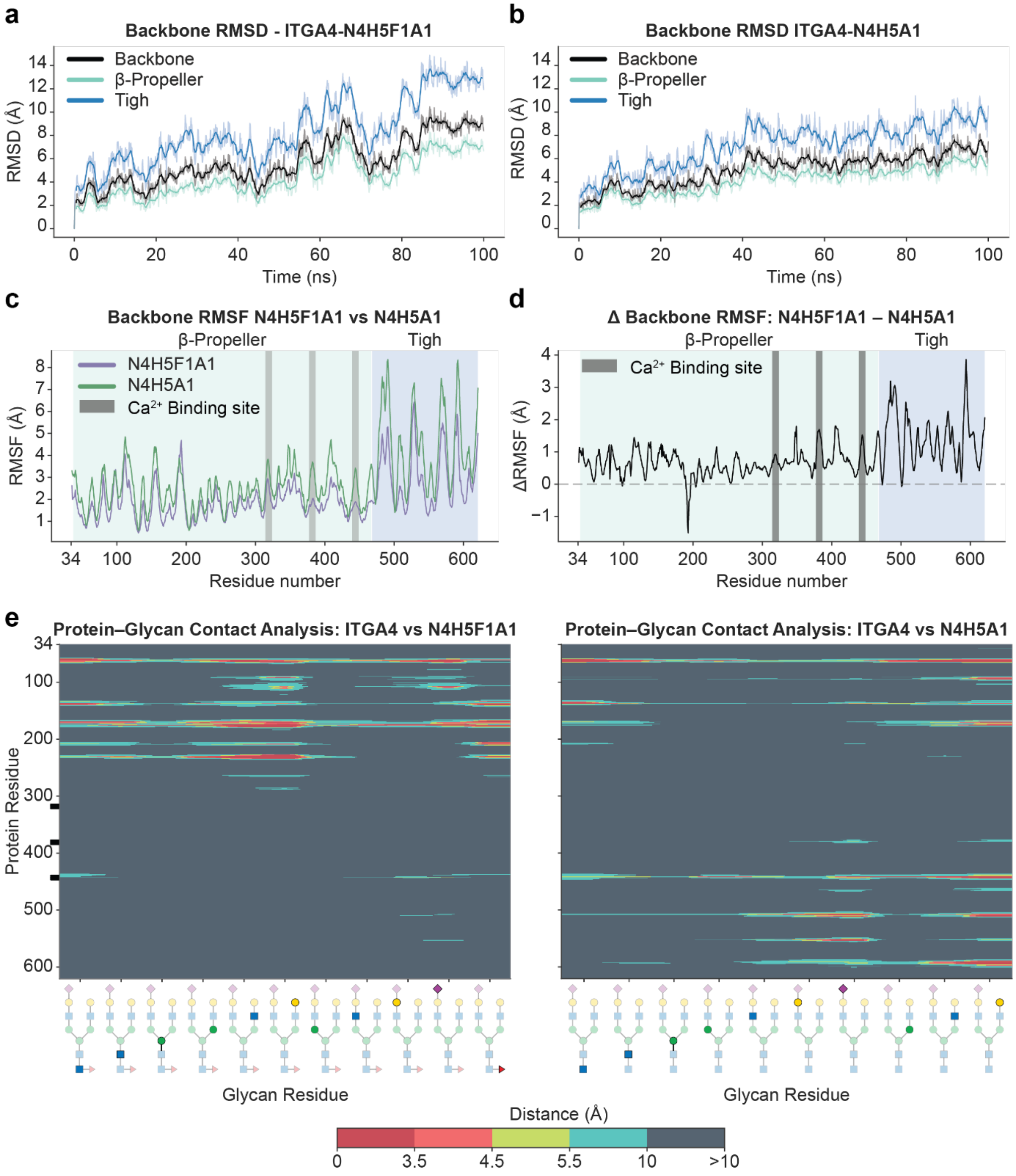


**Extended Data Fig. 7 Molecular dynamics simulations reveal that loss of fucosylation reduces ITGA4 conformational flexibility. (a)** Backbone RMSD over 100 ns MD simulation for fucosylated ITGA4 (N4H5F1A1). RMSD traces shown for overall backbone (black), β-Propeller domain (cyan), and Thigh domain (blue). Dark lines represent 1 ns rolling averages. The Thigh domain exhibits higher structural deviation compared to other regions. **(b)** Backbone RMSD over 100 ns MD simulation for afucosylated ITGA4 (N4H5A1). RMSD values are generally lower and more stable compared to fucosylated ITGA4, particularly in the Thigh domain, indicating reduced overall conformational flexibility. **(c)** Per-residue backbone RMSF comparison between fucosylated (N4H5F1A1, purple) and afucosylated (N4H5A1, green) ITGA4. Gray shaded regions indicate $Ca^{2+}$ binding site loops. The β-Propeller domain (residues ~34-400) and Thigh domain (residues ~478-599) are labeled. Fucosylated ITGA4 shows consistently higher RMSF values, particularly in the Thigh domain and $Ca^{2+}$ binding regions. **(d)** Difference in backbone RMSF (ΔRMSF = fucosylated - afucosylated) across ITGA4 residues. Positive values indicate regions with greater flexibility in fucosylated ITGA4. $Ca^{2+}$

binding site regions and the Thigh domain show substantial increases in flexibility (ΔRMSF 1-4 Å) in the fucosylated form, consistent with enhanced conformational mobility required for integrin activation. **(e)** Protein-glycan contact frequency maps over the MD simulation. Left: Fucosylated ITGA4 (N4H5F1A1). Right: Afucosylated ITGA4 (N4H5A1). Y-axis shows protein residue number, x-axis shows glycan residue (schematic structures shown below, with the solid glycan moiety indicating the respective residue being measured while other moieties are shown with reduced opacity). Color intensity represents contact distance (Å), with warmer colors indicating closer contacts (<5 Å). Fucosylated glycans maintain contacts primarily near the glycosylation site (N138), while afucosylated glycans establish extensive contacts with residues throughout the β-Propeller and Thigh domains, including $Ca^{2+}$ binding regions (residues ~200-250, 377-385, 439-447).
